# The Identification Markers of activated myofibroblast subsets in the Human Lung Fibrosis Ecosystem via integrated omics Analysis

**DOI:** 10.1101/2024.07.21.604481

**Authors:** Ying Zheng, Zhihong Song, Shifeng Li, Bin Cao, Hongping Wu

## Abstract

**Background:** The aberrant remodeling of the extracellular matrix (ECM) is closely associated with lung fibrosis. However, the mechanisms underlying ECM remodeling in pulmonary fibrosis (PF) remain unclear. The advent of single-cell RNA sequencing (scRNA-seq) has provided valuable insights into the diverse phenotypic and functional characteristics of human PF. Nevertheless, the dynamic of ECM remodeling in terms of ECM synthesizing and the potential activating markers of myofibroblasts in the human PF microenvironment still needs to be investigated.

**Methods:** We performed integrative scRNA-seq analyses on high-fidelity PF data from a public platform by filtering out the low-quality counts and doublets using two doublet prediction methods. Next, we investigated the dynamic of the ECM signature in diverse cells in PF and screened the potential markers of myofibroblasts via fitting a successful polynomial regression model. Finally, the markers of activated myofibroblasts were identified using bulk RNA-seq of pulmonary tissue.

**Results:** First, we depicted the pathogenic landscape and demonstrated the heterogeneity of ECM in PF by integratively analyzing single-cell RNA-seq data, and we hypothesized that myofibroblasts played a significant role in ECM formation. Second, our results successfully displayed the biological dynamic changes of ECM and investigated the 73 positive correlated genes of myofibroblasts in PF via a polynomial regression model. Then, the bulk RNA-seq results further identified eight new activating markers of myofibroblasts, such as MFAP2, MXRA5, and LRRC17 via transcriptomic signature, correlation and ROC scores. Finally, the results of cell-cell interaction indicated that myeloid cells may be involved in regulating ECM remodeling through proliferation mediated by myofibroblasts that secrete POSTN, suggesting that ECM remodeling in PF is a complex and multi-participated process.

**Conclusions:** In summary, we provided insights into the contributions of ECM in human PF by integrative analysis and highlighted potential clinical utilities of myofibroblast subsets as therapeutic targets.

## 1 INTRODUCTION

Nearly 20% of various forms of interstitial lung disease (ILD) of both known and unknown causes could induce varying degrees of pulmonary fibrosis and respiratory malfunction[1]. Pulmonary fibrosis (PF) is characterized by the deposition of collagen and other extracellular matrix (ECM) molecules in the lung, leading to high levels of morbidity and mortality worldwide[2]. Persistent infections, autoimmune reactions, allergic responses, chemical insults, radiation, and tissue injury have been reported as causes of PF[3]. PF can impair alveolar gas exchange and reduce lung compliance even leading to progressive respiratory failure and eventual death[4]. The complex pathogenesis of PF involves multiple cell types, such as injury and loss of alveolar epithelial cells[3, 5], expansion of reparative basal stem cells[6], transitional profibrotic macrophage localized to the fibrotic niche[7], and increased infiltrating PD1^+^CD4^+^ T cells[8]. Myofibroblasts were considered the primary cell type producing collagen in the process of PF[5].

The mechanisms of ECM remodeling in PF have been investigated in past years. Myofibroblasts are activated by a variety of effectors such as transforming growth factor β (TGFβ), angiogenic factors (VEGF), platelet-derived growth factor (PDGF), and WNT signals[3, 7, 8]. Then, the excessive accumulation of ECM proteins (mainly fibrillar collagens) produced by myofibroblasts often disrupts the lung structure and leads to lung dysfunction[9, 10]. At the same time, some cytokines and chemokines like interleukin 13 (IL-13) and C-C motif chemokine ligand 2 (CCL2) have been identified as important regulators of PF[11, 12]. However, hepatocyte growth factor (HGF) could directly and indirectly inhibit ECM remodeling by inducing the survival or proliferation of pulmonary epithelial and endothelial cells while reducing the accumulated myofibroblasts[13]. Last but not least, the degradation of ECM has also been mediated by a series of proteases, such as matrix metallopeptidase 2 (MMP2) and TIMP metallopeptidase inhibitor 2 (TIMP2)[14, 15]. The imbalance of synthesis and degradation of ECM can lead to fibrosis in the late stage.

In recent years, many candidate targets have been identified[16]. However, the pathogenesis of PF is so complicated that it has not been fully defined, and effective targets for treatments are still lacking[17]. The development of single-cell RNA sequencing (scRNA-seq) provides high-resolution insights into the heterogeneity of PF[4, 5, 7, 18–21] and displays the transcriptomic signature and communication of different cell types in FME. However, a study regarding scRNA-seq application for systematically evaluating the contribution of various cell types to ECM regulation in PF is still absent.

Given this, we described the pathogenic landscape and displayed the dynamic changes of different cell types in PF by integrative analyzing scRNA-seq data from three published datasets consisting of eight idiopathic pulmonary fibrosis (IPF) patients, six systemic sclerosis-associated interstitial lung disease (SSc-ILD) patients and 14 donor samples. In this work, the contribution of multiple cell types to ECM regulation and their interactions were systematically evaluated. Our results indicated that myofibroblasts might be responsible for the overexpression of collagens and interaction with other cells through POSTN and some other proteins encoded by new candidate genes identified from this subtype in PF. These findings will provide a reference, helping us understand PF more comprehensively, and identifying potential targets for fibrosis therapies.

## 2 RESULTS

### 2.1 An overview of the single-cell atlas in human PF

To determine the resident human lung cell types that secrete and regulate ECM during homeostasis and PF, we first compiled scRNA-seq data from three published datasets. Then we generated a single-cell map of the lungs from different forms of PF and provided clinical features associated with the samples (Table s01). After filtering and removing the potential doublets from these data (Table s01), we obtained single-cell transcriptomes from 163,914 cells and clustered them into six main cell populations: mesenchymal cells (Mes), endothelial cells (Endo), alveolar epithelial cells (AEC), myeloid cells (Mye), natural killer cells & T cells (NK & T), B cells & Plasma cells (B & PC), which were further identified into 22 clusters by relative specific markers (Fig. 1b, d and Fig. s1a, b). Next, we analyzed the proportions of these clusters in PF and observed an increased proportion of mesenchymal cells in the IPF and SSC-ILD groups (Fig. 1e and Fig. s1c). Additionally, mesenchymal cells were the dominant cell type in the UMAP area of the IPF group compared to healthy donors (Fig. 1c, g).

**Fig. 1.**
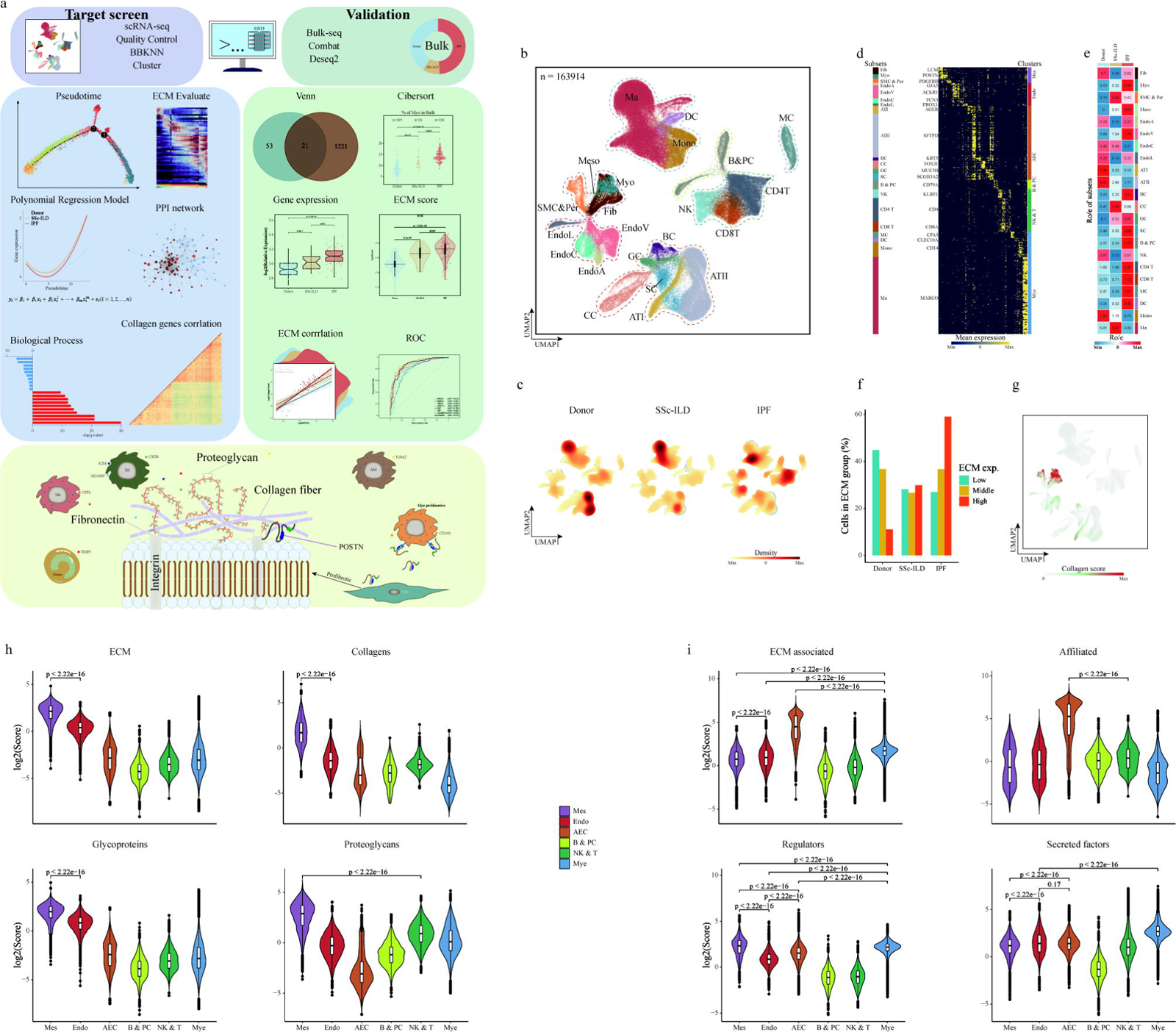
Study design and integrative scRNA atlas of PF. a, Workflow of study. b, UMAP embedding of 163,914 single cells from 28 human lungs. Labels refer to 22 clusters identified. ATI, alveolar epithelial type 1 cell; ATII alveolar epithelial type 2 cell; BC, basal cell; B&PC, B cell& plasma cell; CC, ciliated cell; CD4T, CD4(+) T cell; CD8T, CD8(+) T cell; DC, dendritic cell; EndoA, Endothelial artery cell; EndoC, Endothelial capillary cells; EndoL, Endothelial lymphatic cell; EndoV, Endothelial vein cell; Fib, fibroblast; GC, goblet cell; Ma, macrophage; MC, mast cell; Meso, mesothelial cell; Mono, monocyte; Myo, myofibroblast; NK, natural killer cell; SC, secretory cell; SMC& Per, smooth muscle cells &pericyte. c, Density plots of cells in Donor, SSc-ILD and IPF groups. d, Heatmap of five signature variable genes in each cluster. Each column is the average expression of all cells in a cluster. e, The group prevalence of each cluster is estimated by the Ro/e score. f, All cells clustering by ECM score stratified by groups. g. Feature plot of distribution for collagen score. h, Scores of ECM, collagens, glycoproteins, and proteoglycans for six main cell types. i, Score of ECM-associated genes, ECM-affiliated genes, regulators and secreted factors for six main cell types.

To further understand the main cell types that contribute to the production and regulation of ECM in IPF and SSC-ILD, we established a PF ECM expression score that included collagens, glycoproteins, and proteoglycans, and ECM-associated expression score that included ECM-affiliated proteins, ECM regulators and secreted factors[22–24]. We confirmed that compared with healthy donors, the IPF and SSC-ILD groups had higher ECM expression scores. In contrast, ECM-associated patterns were higher in the donor group, indicating that the expression abundance of ECM genes in IPF and SSC-ILD was higher, while the biological processes associated with ECM were more complicated (Fig. s1d). In addition, ECM scores indicated a clear shift towards high ECM-expressing cells in IPF and SSC-ILD (Fig. 1f). We next analyzed the ECM expression score and ECM-associated expression score in six main cell types and found that Mes had a significantly higher score in ECM expression score while AEC had a higher score in ECM associated expression score due to a higher ECM affiliated score (Fig. 1h, i).

To dissect the reasons for higher ECM-associated expression scores in AEC, we examined the expression abundance of surface protein in main cell types and found that genes encoding surface protein of AEC were a large proportion of ECM-affiliated genes (Fig. s1e). Next, we analyzed ECM expression score and ECM-associated expression score in main cell types and subtypes of endothelial cells as well as AEC (Fig. s1g-i), unstratified and stratified by groups respectively (Fig. s2a-d). The Mes and Mye populations had a relatively higher regulator score while the secreted factors related to ECM were higher expressed in Mye (Fig. 1i). Taken together, these results suggested that a single-cell snapshot of human PF was captured and potential relationships of cell types involved in the biological processes of ECM were also systematically evaluated by integrative analysis.

### 2.2 The contribution of ECM profiles in mesenchymal subtypes for human PF

As aforementioned, excessive accumulation of ECM by mesenchymal cells could lead to fibrosis, especially for fibroblast and myofibroblast subsets[25]. These findings suggested that mesenchymal cells were highly heterogeneous. To further identify the cell subtypes of Mes that mainly contribute to the production and regulation of ECM in PF, we thus profiled 9,621 Mes and clustered them into eight subtypes based on relatively specific markers (Fig. 2a, b and Fig. s3a). Our results showed that POSTN and CTHRC1 genes were highly expressed in myofibroblasts, consistent with a previous study[26]. Furthermore, the ratios of myofibroblasts increased in IPF and SSc-ILD than that in the donor group (Fig. 2c). We additionally found that the proportion of myofibroblasts in Mes populations was higher in IPF and SSc-ILD than that in the donor group and the difference was statistically significant among IPF, SSc-ILD and donor groups (Fig. 2d), demonstrating that myofibroblasts played an important role in causing PF. Next, we studied the ECM expression score and ECM-associated expression score of mesenchymal subtypes. The ECM expression score in IPF and SSc-ILD and ECM-associated expression score in IPF were higher than that in the donor group (Fig. 2e). The ECM expression score including glycoproteins and proteoglycans scores were highest in Fib3 while collagens score was specifically higher in myofibroblasts, indicating myofibroblasts played a relatively key role in producing collagen of ECM. In terms of ECM-associated expression score, Fib3 had a higher secrete factor score than other subtypes whereas myofibroblasts had a higher regulators score, suggesting that different subtypes of Mes subsets played different and complicated roles in contributing to ECM formation and needed to be further investigated (Fig. s3b, c).

**Fig. 2.**
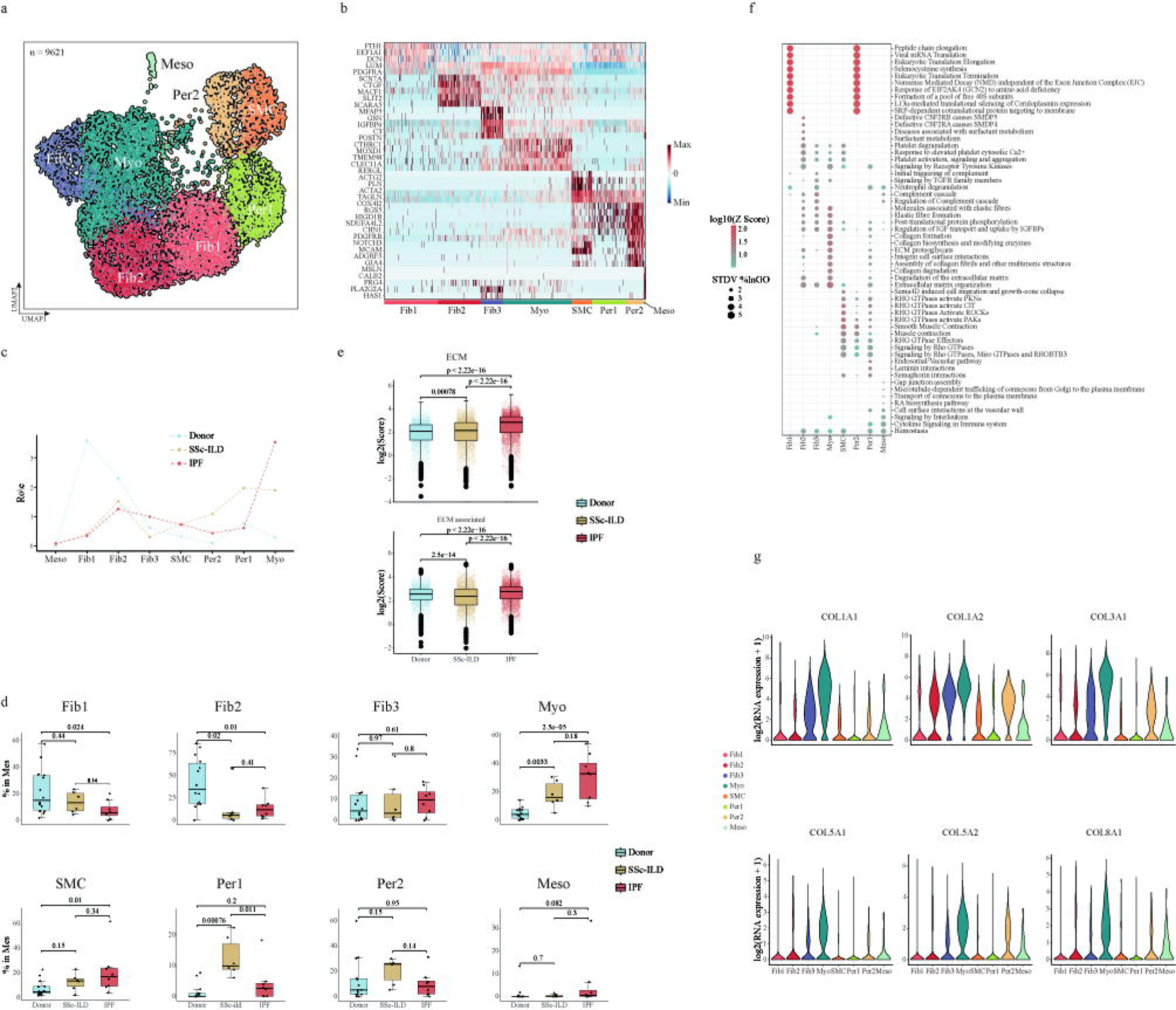
The contribution of ECM profiles in mesenchymal subsets to PF. a, UMAP embedding of 9,621 single cells in mesenchymal subsets. Labels refer to 8 subtypes identified (see Fig.1). b, Heatmap of relative expression of marker genes for mesenchymal subsets. c, Group preference of each mesenchymal subset measured by Ro/e. d, Boxplots of ratios of each mesenchymal subset in 3 groups, respectively. e, Score of ECM and ECM-associated genes stratified by groups in mesenchymal cells. f, Enrichment analysis for Gene Ontology biological processes for each mesenchymal subset. g, Expression of selected collagen-associated genes in mesenchymal subsets.

To investigate the function of different mesenchymal subtypes, we next analyzed the pathways related to ECM in PF of mesenchymal subsets (Table s02). Our analysis uncovered that the transcriptome of myofibroblasts was relatively specifically enriched in collagen formation, collagen biosynthesis and modifying enzymes, assembly of collagen fibrils and other multimeric structures, and collagen degradation (Fig. 2f) (Table s01). We further discovered that Myofibroblasts may play a profibrotic role by expressing more collagens, mainly in the disease. Excepting fibrosis marker such as COL1A1, COL1A2 and COL3A1, we also found COL5A1, COL5A2, COL6A1, COL6A2, COL6A3, COL8A1 highly expressed in myofibroblasts (Fig. 2g and Fig. s3d). We also showed gene expression patterns of ECM expression score and ECM-associated expression score in mesenchymal subtypes (Fig. s3d-h), unstratified and stratified by groups (Fig. s4a-e). Of note, myofibroblasts could have a higher abundance of collagens and regulators in disease (e.g., COL3A1, COL1A1, CTSB, SERPINH1) while Fib3 was more active in disease groups than control from the score results of glycoproteins (e.g., FBLN1, EFEMP1), proteoglycan (e.g., DCN, OGN), and secreted factors (e.g., S100A4, SFRP2) (Fig. s4a-e). Thus, many mesenchymal subtypes, mainly Fib3 and myofibroblasts, regulate the ECM production in human PF, with different patterns.

### 2.3 Trajectory analysis of mesenchymal cells and the dynamic of driving genes for human PF

To decode the transcriptional dynamics of mesenchymal cells, we used VECTOR[27] to describe the trajectory and we observed a clear directional flow from Fib2 to other mesenchymal subtypes (Fig. 3a). Consistent with this result, pseudotime trajectory analysis[28] demonstrated differentiation trajectory of Fib2 to myofibroblasts as well as Fib3 and SMC (Fig. 3b and Fig. s5a-d), which further suggesting that these cell subtypes were most likely to have differentiated from Fib2. We then modeled gene expression along the lineages of mesenchymal cells to identify their gene sets with specific expression patterns in each trajectory (Fig. 3c). Notably, we confirmed that myofibroblasts marked by POSTN and CTHRC1 were at the terminal of one direction (Fig. 3c). We then fitted the ECM score and related genes expression profile to pseudotime using polynomial regression model to investigate the dynamics of ECM patterns and screen candidate genes (Fig. 3d, e and Fig. s3d, e). Collagens score and ECM-associated score also increased higher in IPF and SSC-ILD along pseudotime than that in the donor group (Fig. 3d and Fig. s5d, e), indicating that cell subtypes of terminal state possibly played more significant roles in PF. We further inspected the collagens genes (e.g., COL1A1, COL1A2, COL3A1, COL5A1) expression signature along pseudotime (Fig. 3e and Fig. s3i). Our results showed that all these collagens-associated genes increased along pseudotime in myofibroblasts, suggesting myofibroblasts in mesenchymal cells contributed to collagens formation in ECM, consistent with the previous results (Fig. 2g). And the candidate genes may play similar functions like these collagens associated genes. In summary, our data indicated that myofibroblasts in mesenchymal cells differentiated from Fib2 and might play key roles in collagen formation in human PF.

**Fig. 3:**
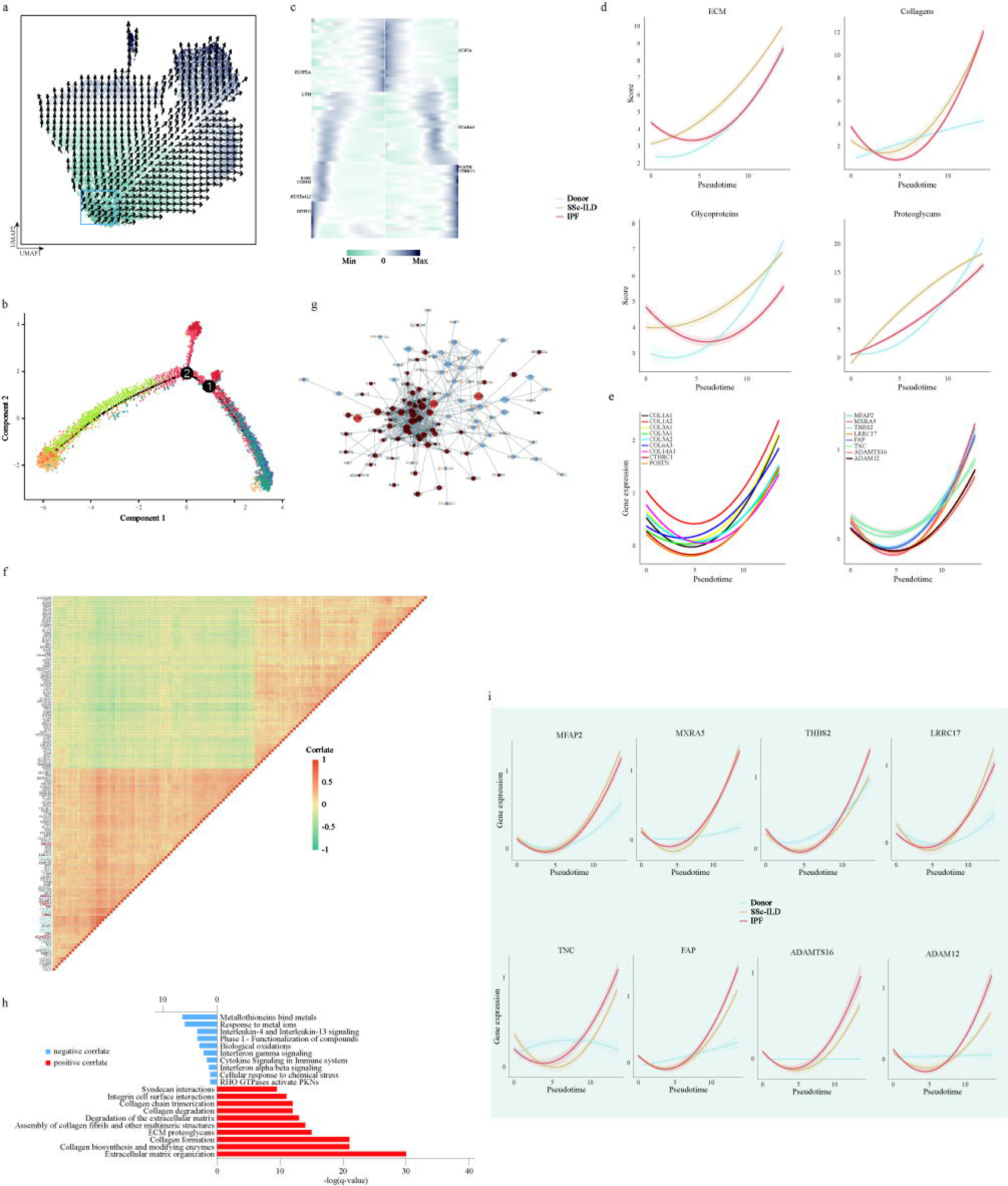
Trajectory analysis of mesenchymal cells and identification of potential therapeutic targets for human PF. a, Pseudotime ordering of mesenchymal cells during PF transformation shown by color (from: Cadet Blue most undifferentiating state to: Dark Byzantine Blue mature state) based on VECTOR. The blue rectangle highlights the area of root cells in UMAP. b, Trajectory analysis of mesenchymal cells. Each dot corresponds to one cell, colored according to its cell subtypes. c, Heatmap of selected gene expression pattern along the pseudotime in two directions in mesenchymal cells. d, Scores of ECM, collagens, proteoglycans and glycoproteins in myeloid cells stratified by groups plotted along the pseudotime. e, Collagen-related genes expression and candidate key genes expression in mesenchymal cells plotted along the pseudotime. f, The correlation analysis of the relative expression level of common genes (see Fig.s5) in myofibroblasts. Candidate key genes are highlighted in red. g, The protein-protein interaction (PPI) networks of these common genes based on STRING. h, Biological processes enrichment for positive and negative genes from these common genes using -log10(Pvalue), respectively. i, Candidate key gene expression in myofibroblasts stratified by groups plotted along the pseudotime.

Since myofibroblasts exhibited a critical function in ECM remodeling, we next investigated the potential dynamic mechanism of PF by gene-related expression modules in myofibroblasts. A total of 137 genes, such as POSTN, CTHRC1, MFAP2 and MXRA5, were acquired from the top 200 genes of positive and negative correlate about COL1A1, COL1A2, COL3A1, respectively (Fig. 3f and Fig. s5g). Further results showed that these positive correlate genes were almost highly expressed in myofibroblasts (Fig. s5g, h) and their biological process mainly were involved in ECM remodelings such as extracellular matrix organization, collagen biosynthesis and modifying enzymes, collagen formation (Fig. 3h). These genes which were higher expressed in IPF and SSc-ILD than donor group were associated with fibrosis pathway and they have strong interaction with each other according to string results (Fig. 3g). The expression of these genes (such as MFAP2, MXRA5 and THBS2) also were higher expressing with pseudotime similar with the patterns of collagen genes (Fig. 3i). Our data indicated above these genes like MFAP2 could be the key gene in ECM remodeling since the strikingly similar pattern with two known ECM regulating genes (POSTN, CTHRC1) in the previous study (Fig. 3i, and Fig. s5i). Together, we found some candidate activating genes by analyzing gene expression modules in myofibroblasts that might play critical roles in human PF.

### 2.4 Identification of the activating markers of myofibroblasts as potential therapeutic targets for human PF

To further validate the above results, we collected the 257 RNA expression profiles by high throughput sequencing from seven public data sets of human lung tissue and biopsy (Table s03, s04). The 1242 high-expression genes were obtained by comparing PF (including 26 SSc-ILD and 126 IPF patients) with 105 donors by Deseq2 (Fig. 4a)[29]. The Venn results indicated that 22 common genes both significantly highly expressed in myofibroblasts correlated gene subset and differentially expressed genes (DEGs) of bulk results (Fig. 4b). The expression results of scRNA altas further identified the 17 genes (COL1A1, COL1A2, COL3A1, COL5A1, COL5A2, COL6A3, COL14A1, CTHRC1, POSTN, MFAP2, MXRA5, THBS2, LRRC17, FAP, TNC, ADAMTS16, and ADAM12) were relatively high expressed in myofibroblasts compared with other cell types (Fig. s6a). We also estimated the ratio of myofibroblasts in bulk data and found that the ratios of myofibroblasts also clearly increased in the SSc-ILD and IPF groups compared to the donor group. Notably, there was even a significant difference between SSc-ILD and IPF (Fig. 4c, Fig. s7a). Finally, these 17 genes were more expressed in SSc-ILD and IPF groups than in donors, and the IPF group was significantly more abundant than the SSc-ILD group in these genes (Fig. 4d, Fig. s7b). There were no significant differences among the 73 residual genes from myofibroblast subsets because of the heterogeneity of PF (Fig. s8a).

**Fig. 4.**
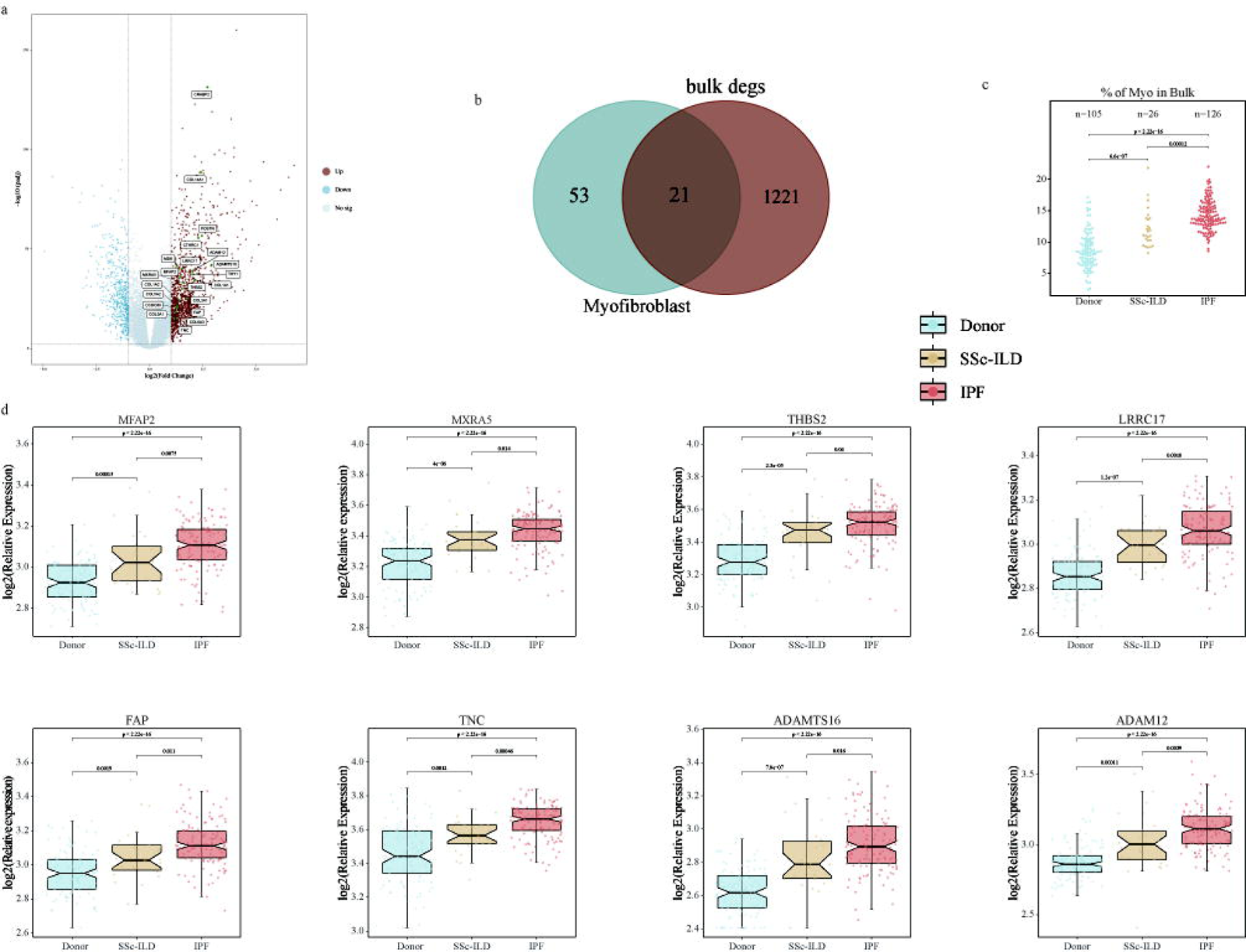
Different expressions of activating genes in myofibroblast in Bulk RNA-seq. a, Volcano plot showed the DEGs of the PF group compared with the donors. b, Venn plot showed the common genes of myofibroblasts and bulk RNA-seq. c, Beeswarm plot showed the estimated ratios of the myofibroblast subset in donor, SSc-ILD and IPF groups. d, Boxplot showed the relative expression of eight newly identified activating genes in different groups.

Except for collagen genes, our ECM score results suggested the SSc-ILD and IPF have high significant expressions of glycoproteins and proteoglycans than the donor and IPF groups notably exhibited the highest ECM scores (Fig. 5a). ECM-associated scores increased in both the SSc-ILD and IPF groups, especially ECM regulators and ECM secreted factors. However, ECM-affiliated scores significantly decreased in the SSc-ILD and IPF groups, which suggested that major functional cells, such as epithelial cell subsets, have been severely damaged, according to our single-cell RNA sequencing results (Figure s8b). We further investigated the potential relationships of the above genes with Collagen levels. We showed eight genes such as MFAP2, MXRA5, THBS2, and LRRC17 had stronger positive correlations with collagens like COL1A1, COL1A2, COL3A1 (Fig. 5b, Fig. s8c). MFAP2 and MXRA5 have even higher levels in the PF groups than POSTN and CTHRC1, the functions of which have recently been proven in fibrosis (Fig. 5b, Fig. s8c). Finally, the ROC results of these 17 genes also suggested that the ROC scores of eight new genes were nearly 80% and showed a similar pattern to that of collagen genes. (Fig. 5c, Fig. s8d).

**Fig. 5.**
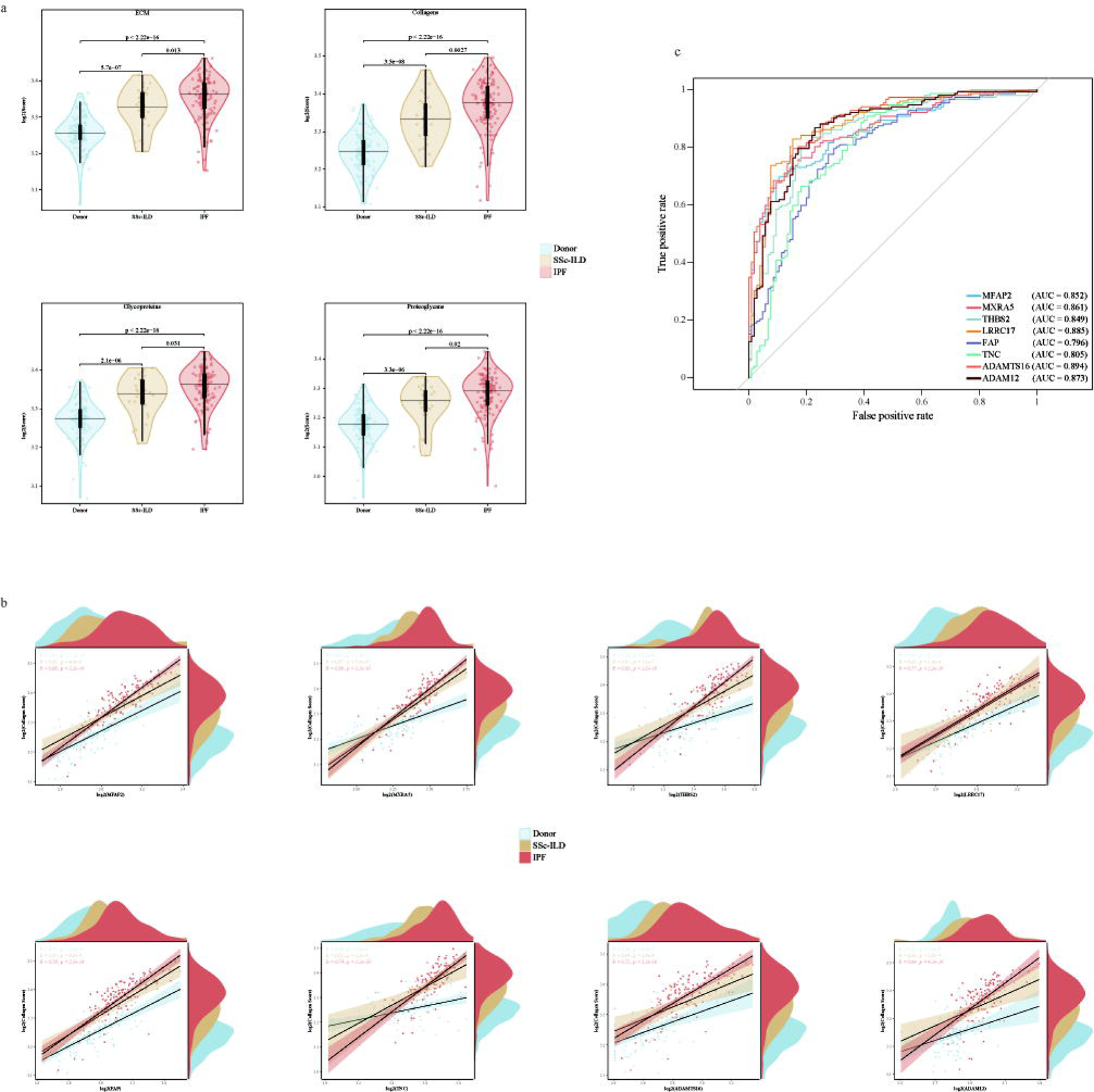
Identification of activating genes in Bulk RNA-seq. a, The violin plot showed the scores of ECM, collagens, glycoproteins and proteoglycans for the donor, SSc-ILD and IPF groups. b, The scatter plot showed a strongly positive correlation between the eight new genes and the collagen score in the SSc-ILC and IPF groups. c, The ROC plot showed the favorable results of these eight new genes in the PF and donor groups.

### 2.5 Heterogeneity of myeloid cells and lymphoid cells involved in the regulation of ECM

The myeloid cells may regulate the synthesis and degradation of ECM from the results described previously[2], we re-clustered 62,859 cells and then identified eight distinct subsets including alveolar macrophage (AM), dendritic cell (DC), interstitial macrophage (IM), macrophage (Ma), mast cell (MC), monocyte (Mono), Myeloid proliferative cell (Mye proliferative) using relative specific markers (Fig. 6a, c, d). Next, we studied the proportions of these subsets in the IPF, SSC-ILD, and donor group. The results revealed that compared to the donor group, the proportions of MC, DC and IM increased in IPF, while Ma elevated in SSc-ILD (Fig. 6b). IM and Ma, which derived from monocyte, obviously could be involved in ECM and collagen-associated biology process from the GO enrich terms and these results indicated that the regulation of ECM by myeloid cells was complex (Fig. 6e) (Table s05).

**Fig. 6.**
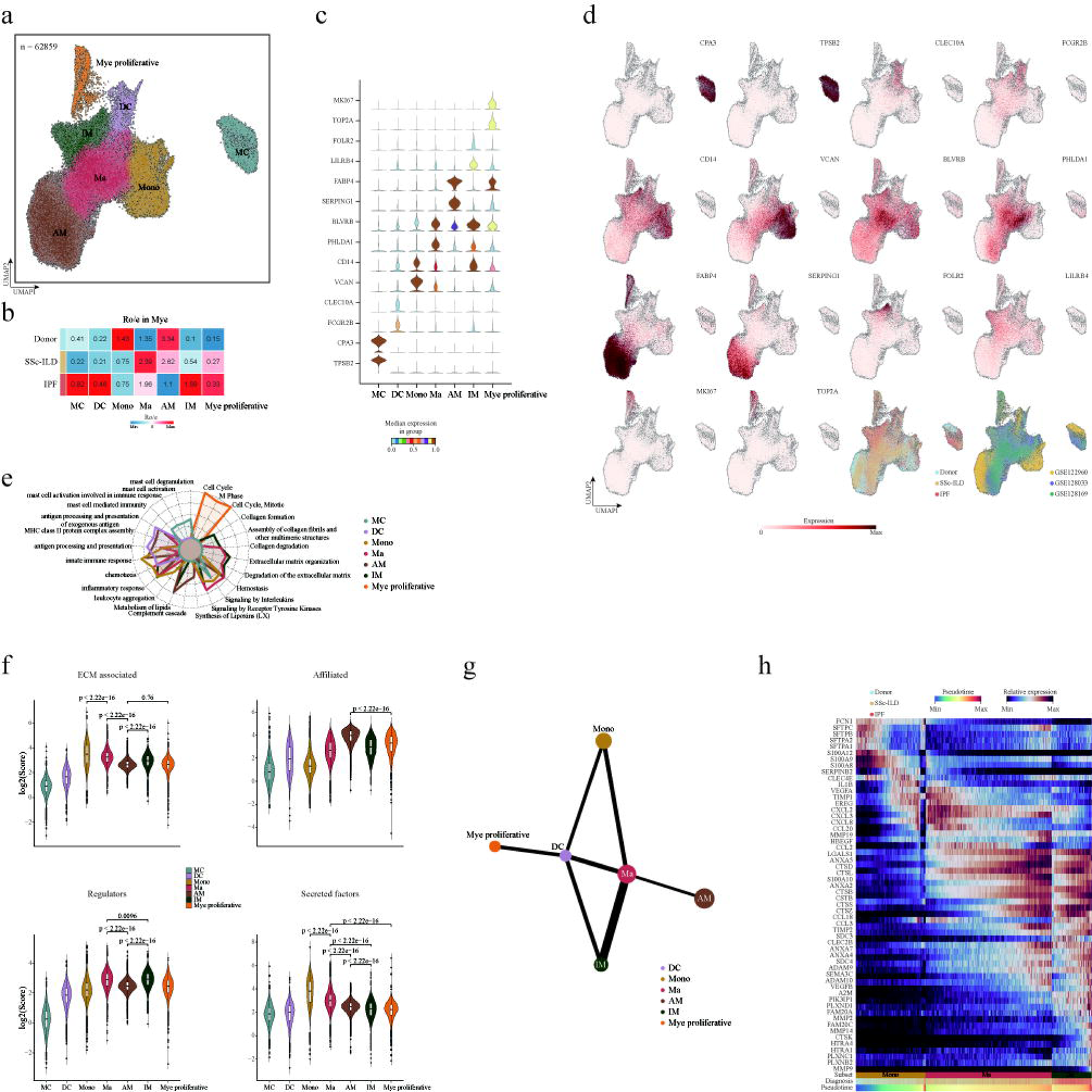
Heterogeneity of myeloid cells involved in the regulation of ECM. a, UMAP embedding of 62,859 single cells in myeloid subsets. Labels refer to 7 subsets identified.AM, alveolar macrophage; DC, dendritic cell; IM, interstitial macrophage; Ma, macrophage; MC, mast cell; Mono, monocyte; Mye proliferative, Myeloid proliferative cell. b, The group prevalence of each myeloid subset is estimated by the Ro/e score. c, Violin plot of expression of classical marker genes of each myeloid subset. d, Expression of classical marker genes of myeloid subsets on the UMAP embedding. Stratification of myeloid subsets by three groups and 3 GEO accessions. e, Radar plot of GO enrichment of highly expressed genes in each myeloid subset. f, Score of ECM-associated genes, ECM-affiliated genes, regulators and secreted factors in myeloid subsets. g, PAGA analysis of main myeloid subsets. Line thickness corresponds to the level of connectivity. h, Heatmap of expression pattern of ECM associated genes in monocyte to interstitial macrophage path along the pseudotime stratified by groups.

Then we evaluated the ECM-associated expression score in each myeloid subset. Monocytes displayed higher ECM-associated secreted factors like S100A8/9 and IL1B while AM contributed to the affiliated genes such as C1QA, C1QB, and C1QC. However, these genes showed limited high expression in the diagnosis. At the same time, we found that the Ma and IM performed higher ECM regulator scores due to the high expression of genes like CTSB, CSTB and TIMP2 (Fig. 6f, Fig. s9a-c). We next used partition-based graph abstraction (PAGA) analysis to examine the connectivity structures between the myeloid subtypes and found that the Ma was derived from the monocytes and they were different from the IM according to the previous studies (Fig. 6g, Fig. s9g). To identify the relationship between subsets and groups, we analyzed gene patterns along pseudotime and found that IM may play a critical role in regulating ECM remodeling in IPF while Ma may regulate ECM remodeling in SSc-ILD (Fig. 6h). Our results further investigated the transcriptomic signature of these ECM-associated genes in different subsets by groups respectively (Fig. s9d-f). All the above findings indicated that the accessibility of dynamic ECM-associated gene motifs across the trajectory was consistent with the sequential differentiation states (Fig. 6h).

To investigate the roles lymphoid cells played in contributing to ECM remodeling in PF further, we analyzed lymphoid cells as described above. We first obtained transcriptomes of 31,698 cells and then identified 12 clusters (B cell; B memory; B naïve; B &PC proliferative; CD4^+^ T memory; CD4^+^ T naïve; Treg; Th2; CD8^+^ T eff; CD8^+^ T memory; NK; PC) based on relative specific markers (Fig. s9h, i). We next examined the proportions of these subsets in the IPF, SSC-ILD and donor groups and the results represented that compared to the donor group, the proportions of B cells (B cell, B naïve, B &PC proliferative) and Treg increased in IPF while they decreased in SSc-ILD, except for B memory which increased in both of them; Conversely, T cells (CD4 T naïve, Th2, CD8T eff, CD8T memory) increased in SSc-ILD while decreased in IPF (Fig. s9j). Finally, the ECM and ECM-associated results of lymphoid indicated that they had limited effect on ECM remodeling in PF (Fig. s9k-l).

### 2.6 Identification of the interactions of main myeloid subsets and myofibroblasts in human PF

Mesenchymal cells are known to interact with other cell types in the lung[5, 17]. Our results showed that interactions among different cell subsets were more abundant in IPF, and SSc-ILD than in the donor group (Fig. 7a and Fig. s10a, c). Consistently, there were higher proportions and numbers of differentially expressed genes (DEGs) in IPF and SSc-ILD compared to the donor group. These DEGs included previously reported fibrosis-associated genes such as PERIOSTIN, TGFβ, and ANNEXIN (Fig. 7b and Fig. s10a, b). We then performed a communication pattern analysis to investigate the communication signaling between myofibroblasts and other cells in PF.

**Fig. 7.**
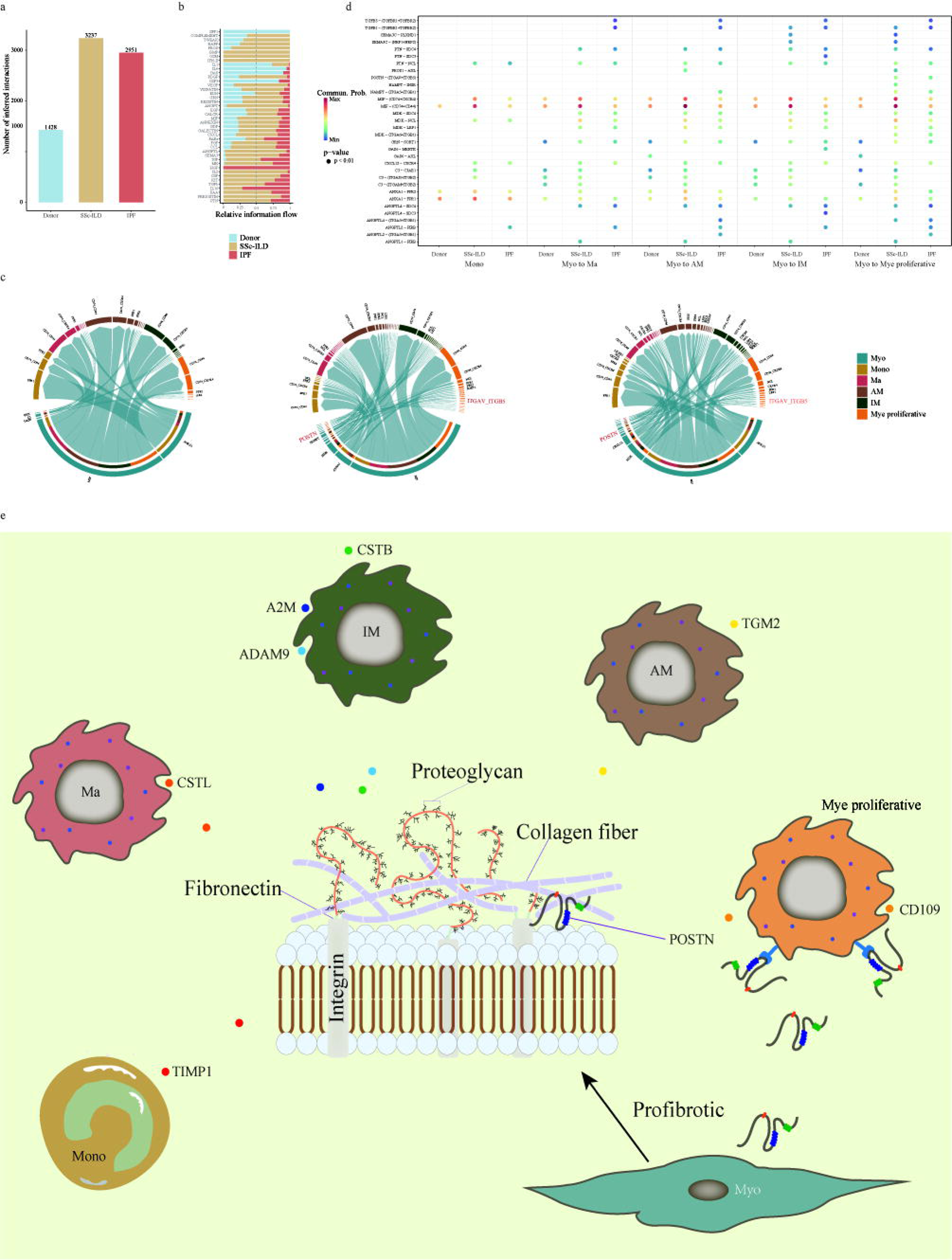
Identification of cell-cell interactions in human PF. a, Number of cell-cell interactions among Donor, SSc-ILD and IPF using CellChat. b, The relative information flow of differentially expressed genes (DEG) for the PF stratified by groups. c, Circos plots of ligand and receptor interactions of myofibroblast subset and main myeloid subsets. d, Comparison of the significant ligand-receptor pairs among Donor, SSc-ILD and IPF. e, Schematic of proposed role myofibroblasts and myeloid subsets in ECM reprograming in PF.

Our results uncovered six distinct patterns for outgoing signaling (Fig. s10d) and incoming signaling (Fig. s10e). The outgoing myofibroblasts signaling in SSc-ILD was characterized by ANGPTL, PIN, PERIOSTIN, KIT and BMP while outgoing myofibroblasts signaling in IPF was represented by FGF, ANGPTL, IGF, PERIOSTIN, HGF, and CSF (Fig. s10d), which was absent in the donor group. On the other hand, the communication patterns of target cells showed that incoming myofibroblasts signaling in SSc-ILD was dominated by PDGF, PARs, PERIOSTIN, EDN, PROS and BMP while incoming myofibroblasts signaling in IPF was enriched by FGF, EDN, PERIOSTIN and PDGF (Fig. s10e).

We next identified ligand-receptor pairs between myofibroblasts and myeloid subsets in IPF, SSc-ILD versus the donor group. Our data showed an increase of MIF ligand-receptor interactions between myofibroblasts and each myeloid subset in SSc-ILD compared to the donor group (Fig. 7d). We also found TGFβ1 ligand-receptor interaction increased only in the IPF group (Fig. 7d), indicating the specific roles of the TGFβ pathway in IPF development. Notably, the POSTN-ITGAV and POSTN-ITGB5 interaction were only increased in myofibroblasts and Mye proliferative in IPF, SSc-ILD compared to the donor group (Fig. 7d). Ligand–receptor circle figures also suggested that fibrosis signaling interaction POSTN between myofibroblasts and Mye proliferative was significantly increased in IPF, SSc-ILD compared to the donor group (Fig. 7c), indicating myeloid cells might be involved in ECM remodeling through proliferation which is mediated by myofibroblasts secreting POSTN (Fig. 7e). Take together, these results suggested that interactions between myofibroblasts and other cells increased in different PF.

## Discussion

In this study, we construct the integrative analysis atlas of ECM remodeling to systematically investigate the contributions of mesenchymal, endothelial, epithelial, B & PC, NK & T and myeloid cells in human PF by analyzing IPF and SSC-ILD data. To overcome the challenges of the doublets and batch from the different scRNA-seq datasets, we first removed the potential doublet cells by using two benchmarking computational doublet detection methods and classical markers used in previous studies on diverse cells. Subsequently, we created a neighbor graph across these data using bbknn to identify the k nearest neighbors for each sample. Our strict strategy was to ensure the high purity and completeness of the PF landscape with minimal impact on downstream proportion comparisons and other bioinformatics analyses. We particularly confirmed that AEC has a higher expression of ECM-affiliated genes compared to other cell subpopulations due to the major biomarkers of ATII, suggesting that it is necessary to screen the ECM gene set to more accurately evaluate ECM remodeling in PF and even fibrosis in other organs. However, it is hard to reconstruct the real PF microenvironment using any computational strategy due to the limited high fidelity and the heterogeneity of datasets from different labs. Therefore, it is necessary to build the transcriptional phenotype of the ECM model and develop a new computational strategy using an interdisciplinary approach to further explore the mechanism of PF, even fibrosis in other organs.

We also provided new insights into the status of distinct mesenchymal subpopulations in PF pathogenesis, especially for myofibroblast subpopulations. Furthermore, we comprehensively assessed the functional diversity of ECM-associated cell types in the PF microenvironment. The myofibroblasts were activated in PF and served as a key source in fibrosis by producing collagen, which may differ from various sources including resident mesenchymal cells, and epithelial and endothelial cells in the process named EMT/EndMT[3]. In our research, myofibroblast subpopulations exhibited substantial PF preferences and the trajectory analysis indicated that the PDGFRA^+^ fibroblast subset (Fib2) as a source could potentially differentiate into myofibroblasts or pericyte subtypes (Per2). Our pseudotime model could interpret the excessive accumulation of ECM components such as collagens. We also further investigated potential dynamic markers driving myofibroblasts to produce pathologic collagens in this study. POSTN, which was positively correlated with the activating markers, was involved in ECM-associated biological processes as previously proven[24]. Recently, some studies have also revealed that CTHRC1 could be a crucial player in fibrosis in cardiac fibroblasts, pulmonary fibroblasts, and even COVID-19[26, 30, 31]. The CTHRC1+ pathological fibroblasts showed a clear profibrotic signature in mouse hearts. This signature emerged in fibrotic lungs, leading to the highest levels of collagens and even contributing to rapidly ensuing PF in COVID-19 as indicated by scRNA-seq[26, 31]. However, the myofibroblast subpopulation was not clearly enough characterized in these works. Our results suggested that CTHRC1^+^ fibroblasts may be the pathological myofibroblast subtype and the CTHRC1 and POSTN could be the potential markers of myofibroblasts.

However, these current findings were still limited to investigate the mechanism and treatment of PF. Our new identification markers such as MFAP2, MXRA5, THBS2, LRRC17, FAP, TNC, ADAMTS16, and ADAM12 could provide new potential insight for the diagnosis and treatment of PF. MFAP2 is a major antigen of elastin-associated microfibrils and recently had been proved to promote HSCs activation through FBN1/TGFβ/Smad3 pathway, of which TGFβ had a decisive player in fibrosis as previously reported[32]. Another study reported that MFAP2 was involved in the invasion and migration of melanoma through EMT and the Wnt/β-Catenin pathway, which also played an important role in fibrosis[33]. At the same time, MFAP2^+^ fibroblasts were strongly negatively correlated with the prognosis and therapeutic resistance of gastric cancer[34]. However, the detailed functions of MFAP2 in fibrosis have received limited attention compared to studies focused on cancer [35]. Additionally, further investigation into LRRC17 is also necessary to explore its potential biological processes and mechanisms in myofibroblasts[36]. The researchers had proved that MXRA5, with anti-inflammatory and anti-fibrotic properties, was a downstream gene regulated by TGFβ1 but not TWEAK in chronic kidney disease[37]. In contrast to the study above, our research demonstrated that MXRA5 had a higher expression in IPF and SSC-ILD compared to the donor group. Other works also reported that MXRA5 was associated with Cystic fibrosis transmembrane regulator (CFTR), IPF and Klinefelter syndrome[38–40].

At the same time, more and more studies have confirmed the functions of THBS2 in diverse fibrosis, even in PF[41–44]. FAP and TNC genes as biomarkers have been proved in liver fibrosis and kidney fibrosis[45, 46]. The relative RNA expression and protein levels of ADAMTS16 elevated in mice with cardiac fibrosis [47]. Some researchers further proved the correlation between ADAMTS16 and CTHRC1 in cancers[48]. ADAM12 has been suggested to be associated with the initiation and progression of dermal fibrosis and systemic sclerosis-associated interstitial lung disease, as indicated by reported serum levels [49]. However, our results confirm that ADAM12 could serve as a potential identification target for activated myofibroblasts based on scRNA-seq landscape analysis and bulk data from lung tissue. These data and our results indicated that the functions and mechanisms of the eight newly identified biomarkers in fibrosis require further investigation in the future. These genes may provide valuable insights for developing novel therapeutic strategies in PF.

Our work also had a systematic evaluation of ECM-associated processes among immune cells, and it was found that the myeloid cells were preferable to influence ECM remodeling compared to lymphatic cells by regulators or secreted factors. Previous studies have indicated that monocytes and monocyte-derived macrophages could play an important role in fibrosis[50, 51]. Interstitial macrophages significantly appeared in the disease and they were involved in ECM remodeling processes such as extracellular matrix organization (A2M, CTSB, CTSS, TIMP2, ADAM9), Collagen degradation (CTSB, ADAM9), and Collagen formation (CTSB, CTSS) (Fig. 6e, Fig. s9e, Fig. s10e). We also found that other myeloid cells, such as alveolar macrophages and monocyte-derived macrophages, could participate in the regulation of ECM by TIMP1, SERPINA1, and CTSL (Fig. s9e, Fig. 10e). Moreover, the cell-cell interactions further demonstrated a significant increase in cellular communication signals within the PF microenvironment compared to the donor group (Fig. 7a, Fig. 7e). Our results also indicated that myofibroblasts may potentially influence the proliferation of myeloid cells via POSTN (Fig. 7d-e).

In summary, our systematic analyses revealed the heterogeneity and complexity of the PF microenvironment and the accurate ECM gene sets needed to be built for human fibrosis in the future.

## Conclusions

We successfully constructed an ECM snapshot of PF via integrated scRNA-seq analysis to investigate the contributions of diverse cells in the lung fibrosis microenvironment. The expression pattern of ECM and trajectory analysis of mesenchymal cells demonstrated that myofibroblasts played a major role in ECM synthesizing signatures. yoBy using a polynomial regression model and the bulk RNA-seq, we successfully identified eight new genes as activating markers of myofibroblasts to diagnose and target therapy in PF. Furthermore, we also investigated the dynamic regulation associated with ECM remodeling of myeloid cells during PF.

## Acknowledgments

We sincerely thank all the fellows, colleagues and collaborators who contributed to the work mentioned in the manuscript. This work was supported by the China Postdoctoral Science Foundation (2020M670360).

## Authors’ contributions

BC and HPW conceived the studies, oversaw the experiments, and wrote the manuscript. YZ, ZHS and SFL performed the studies. All authors read and approved the final manuscript.

## Materials And Methods

### scRNA public datasets acquisition, quality control and visualization

Considering the variety of types and pathogenicity threat of PF, we attempted to integrate the scRNA-seq data of IPF and SSc-ILD for a more comprehensive analysis. Thus, the public scRNA-seq raw count matrices for eight IPF, six SSc-ILD and 14 nonfibrotic control individuals were acquired respectively from GSE122960, GSE128033, and GSE128169[2, 20, 21]. The detailed information about these counts of cells and genes and corresponding clinical data of the patients were in supplement Table S1. The data quality control was assessed via Scanpy (version 1.7.0)[52]. A total of 223291 cells were sorted by filtering out the low-quality counts by cells with less than 200 genes expressed, genes detected in less than three cells and more than 10% mitochondrial-derived genes. At the same time, the doublet cells from the matrix of each patient were evaluated and sorted by Doubletdetection (http://doi.org/10.5281/zenodo.2678041) and Scrublet[53], then these doublets were further identified manually in subsequent cell type analysis by cell expressed specific markers, and removed (Table s01, Fig. s1a). The merged expression 164650 cells matrices were normalized using Scanpy’s pp.normalize_per_cell (counts_per_cell_after = 1e4) and log transformation was used via Scanpy’s pp.log1p() and identified highly-variable genes as protocols. Next, we scaled each gene to unit variance as tutorials, then reduced the dimensionality of the matrices by running principal component analysis (PCA). The neighborhood graph of cells using the PCA representation of the matrices was calculated by n_neighbors = 30 and n_pcs = 15. The neighborhood graph of cells was directly used Leiden[54] to cluster by using the resolution = 1, and these clusters were plotted by uniform manifold approximation and projection (UMAP) finally[52].

### Cell type annotation and batch correction

First, immune cells (PTPRC^+^), including myeloid cells and lymphatic cells, mesenchymal cells (ASPN^+^), endothelial cells (PECAM1^+^/PTPRC^−^), and epithelial cells (EPCAM^+^) were used to split these clusters into six major subgroups. Each subgroup underwent the same dimensionality reduction, clustering, visualization as described above and batch correction using batch balanced k nearest neighbors approach (bbknn). Each cell subpopulation was finally further clustered into cell type and manually annotated by individual markers such as fibroblast (LUM), myofibroblast (POSTN), pericyte (PDGFRB), smooth muscle cell (ACTA2), type I epithelial cells (AGER), type II epithelial cells (SFTPD), basal cell (KRT5), ciliated cell (FOXJ1), goblet cell (MUC5B), secretory cell (SCGB3A2), B cell (CD79A), natural killer cell (KLRF1), T cell (CD3D), CD4^+^T cell (CD4), CD8^+^T cell (CD8A), mast cell (CPA3), dendritic cell (CLEC10A), monocyte (CD14), and macrophage (MARCO) (Fig s1.b)[2, 6, 18, 25]. In our results, 736 erythrocyte cells marked by HBB were not analyzed in the following process. The differentially expressed genes between cell subpopulations were identified by computing a ranking for the highly differential genes in each cluster via pp.highly_variable_genes () and tl.rank_genes_groups(). The Gaussian kernel density estimates of the diagnosis group were embedded by tl.embedding_density () and plotted by pl.embedding_density ().

### Score signature of ECM, pathway, and GO enrichment of cell types and cell status

The signature of various scores about ECM and cell types was summarized based on normalized gene expression data using the same method on cell cycle analysis[55]. We analyzed the ECM score and ECM associated using core matrisome genes and matrisome-associated genes previously described[24]. In this work, the scores of collagens, glycoproteins, proteoglycans, ECM-affiliated genes, ECM-regulators and ECM-secreted factors were also valued using the same algorithm, respectively. The collagen score of each cell in this data was clustered into three groups (High, Middle, and Low) using KMeans imported from sklearn1.2.2. To functionally and annotationally describe the cell subpopulations in PF, we performed the KEGG and Reactome pathway and Gene Ontology (GO) analysis from the metascape[56] by using the top 100 variable genes of each subset via multiple gene lists (Table s01-02). The dot plot about mesenchymal subsets and the radar plot about myeloid subsets were individually plotted with ggplot2 (version 3.4.4) and fmsb (version 0.7.6) in R4.2.0. The results of PPI (Protein-Protein Interaction Networks) about collagens were obtained from String and were visual by Cytoscape (version 3.10.1)[57].

### Trajectory inferences by Monocle2, VECTOR, diffusion maps and PAGA analysis

The inferred developmental directions of mesenchymal cells in UMAP were investigated by VECTOR, and then the trajectory inferences and pseudotime in mesenchymal cells were explored by Monocle2 (version 2.26.0). The ECM scores and the expression profile of target genes were fitted to the pseudotime using a polynomial regression model to investigate the dynamics of ECM patterns and candidate gene screens. To model the potential lineage tree inference about the myeloid cells, the raw counts of target cell subtypes were not preprocessing and only logarithmic to analyze by construction diffusion map by Scanpy function tl. dpt() and the connectivity and distance of the target cell subtypes were performed using PAGA[58] analysis, also in Scanpy (version 1.9.1)[52].

### Cell-cell interaction analysis by cellchat

The normalized gene expression matrix of donor, SSc-ILD, and IPF was individually input in the Cellchat (version 1.6.1)[59] as their tutorials for exploring the cell-cell interactions in different groups. We identified the major signaling changes across the merge results of these three groups as the protocol to investigate cell-cell interactions in PF.

### Bulk public datasets acquisition, quality control, identify DEGs and CIBERSORT

To validate the above results, we collected 257 tissue and/or biopsy samples, including 105 donors, 26 SSc-ILD and 126 IPF samples from GSE231693, GSE52463, GSE83717, GSE92592, GSE213001, GSE199152, and GSE166036. The 15000 common genes of the raw counts with the same Gene ID would merge for the subsequent analysis (detail in Table s03, s04). First, the batch effect was removed by limma::removeBatchEffect() and the normalized and variant stabilized results were utilized to identify DEGs via the DESeq2 (1.38.3) statistical tool[29, 60]. Between the control and PF groups (including SSc-ILD and IPF samples), and genes with a P-adjusted value (adjp) of <.01 and a |log2FoldChange| > 1 were calculated as significant DEGs. The variable gene expression file of PF scRNA would be used as the reference to estimate the ratio changes of bulk RNA seq using CIBERSORT[61].

### Statistical analysis

For most scores and percent of subsets results, comparisons were made using the Wilcox.test using ggsignif (version 0.6.4.9000) in R4.2.0. The ratio of observed to randomly expected cell numbers (Ro/e) was calculated using the chi-square test to examine their diagnosis group preference for each subset. The correlation analysis between collagens genes (COL1A1, COL1A2 and COL3A1) was performed by using Pearson correlation via corrplot (version 0.92) also in R4.2.0. The common genes of three genes of Venn results would also be showed in Venn. The comparisons of bulk DEGs were also made by Wilcox.test using ggsignif as above, and the correlation analysis between target genes and collagens scores was also performed by using Pearson via ggExtra (0.10.0). The ROC scores of target genes in bulk RNA-seq were calculated by pROC (1.18.0).

## Data availability

The public scRNA-seq raw count matrices of this study were acquired from the Gene Expression Omnibus (GEO) with accession number GSE122960, GSE128033, and GSE128169 and the detailed information was in Table s1. The public scRNA-seq raw counts of our study were acquired also from the GEO with accession numbers GSE231693, GSE52463, GSE83717, GSE92592, GSE213001, GSE199152, and GSE166036 and the detail information was in Table s4.

## Code availability

Example scripts to process and analyze code are available at https://github.com/tsinghuavirgil/cjfh_pccm_03_fibrosis. Detailed information will be available from the corresponding author upon reasonable request.

---

**Fig. s1.**
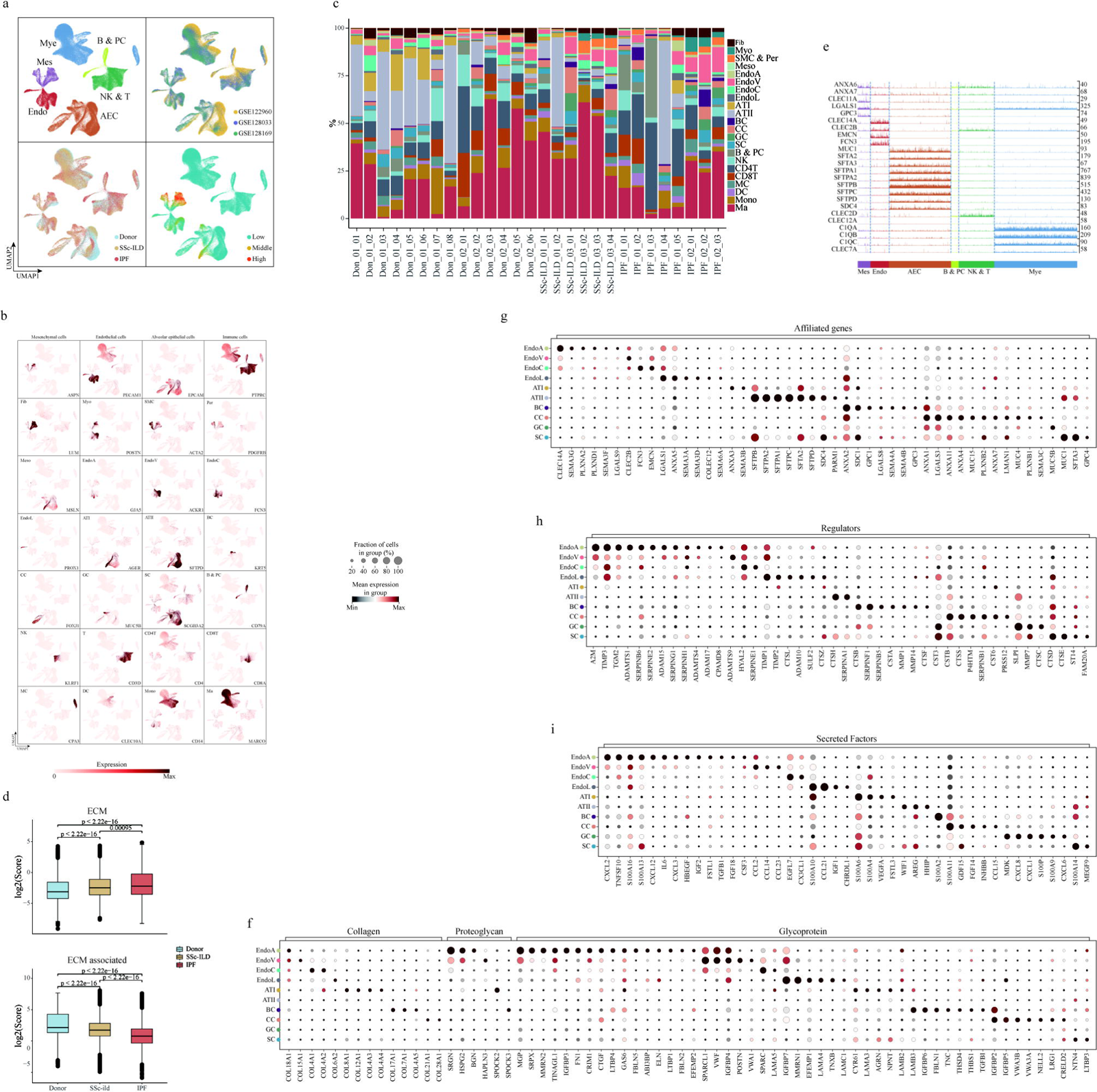
Human lung cell atlas and the expression patterns of ECM and ECM-associated genes. a, Stratification of cells by six main cell types, three GEO accessions, three groups and ECM score. b, Expression of classical marker genes of clusters on the UMAP embedding. c, Frequencies of clusters for individual patients. d, Scores of ECM genes and ECM-associated genes stratified by groups. e, Counts of ECM-affiliated genes in main cell types. f, Dot plot of relative expression of collagens, proteoglycans and glycoproteins associated genes in each subtype of AEC and Endothelial cells. g-i, Dot plot of relative expression of ECM-affiliated genes, regulators and secreted factors in each cell subtype of AEC and Endothelial cells.

**Fig. s2.**
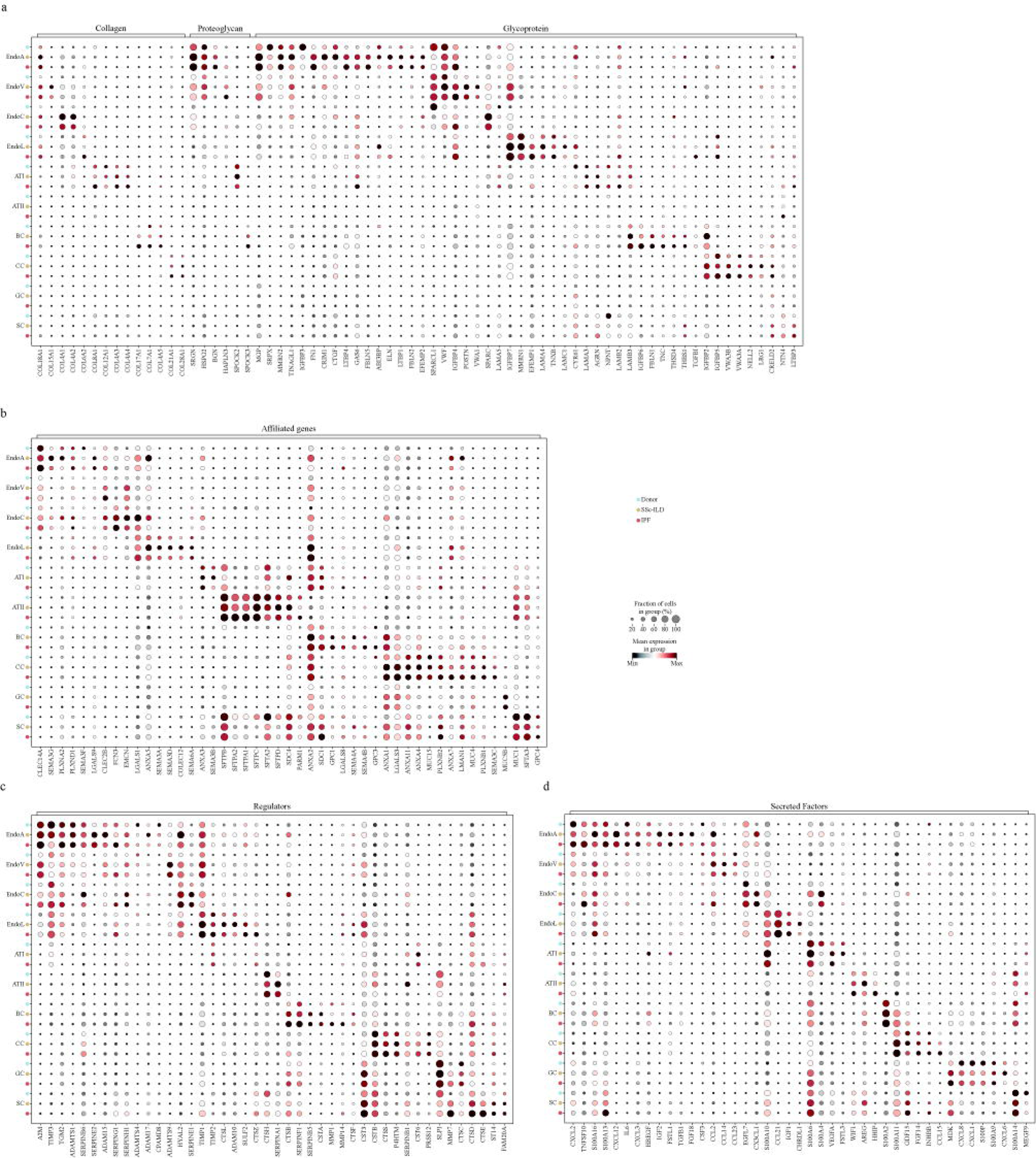
The signature of ECM-related genes in AEC and Endo by group. a-d, Stratification of relative expression of genes in cell subtypes of AEC and Endothelial cells by groups. Dot size and color indicate the fraction of expressing cells and normalized expression levels, respectively.

**Fig. s3.**
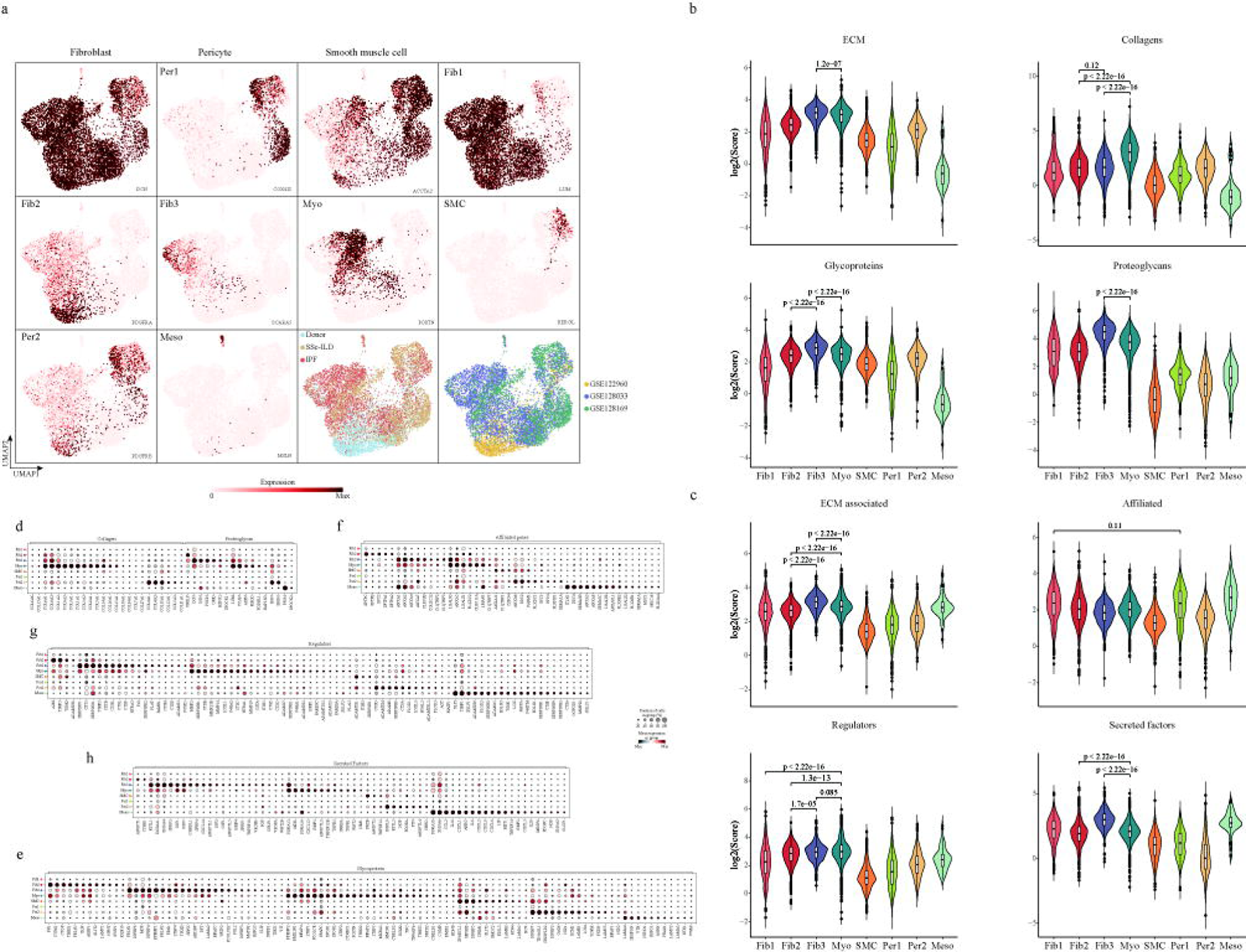
Characteristics of mesenchymal subsets. a, Expression of classical marker genes of mesenchymal subsets on the UMAP embedding and Stratification of mesenchymal subsets by 3 groups and 3 GEO accessions. b, Scores of ECM, collagens, proteoglycans and glycoproteins for each mesenchymal subset. c, Scores of ECM-associated genes, ECM-affiliated genes, regulators and secreted factors for each mesenchymal subset. d-e, Dot plot of relative expression of collagens, proteoglycans and glycoproteins for each mesenchymal subset. f-h, Dot plot of relative expression of ECM-affiliated genes, regulators and secreted factors for each mesenchymal subset.

**Fig. s4.**
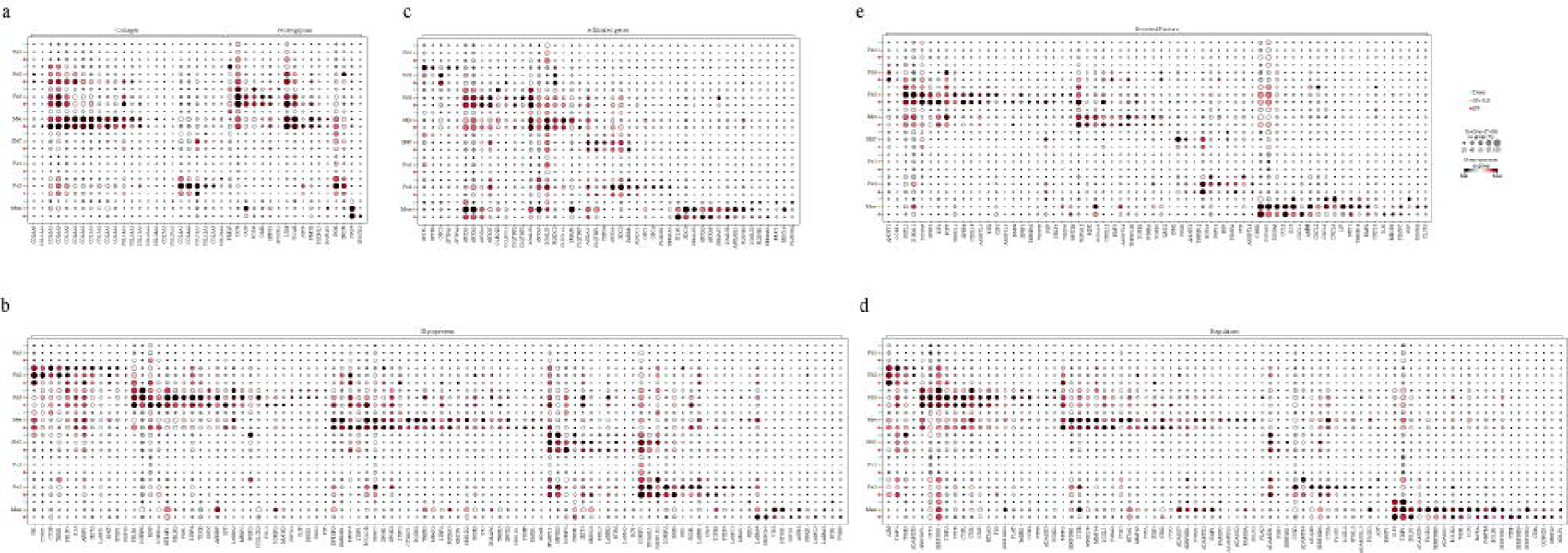
The signature of ECM-related genes in Mes subsets by group. a-e, Stratification of relative expression of genes for mesenchymal subsets by groups.

**Fig. s5.**
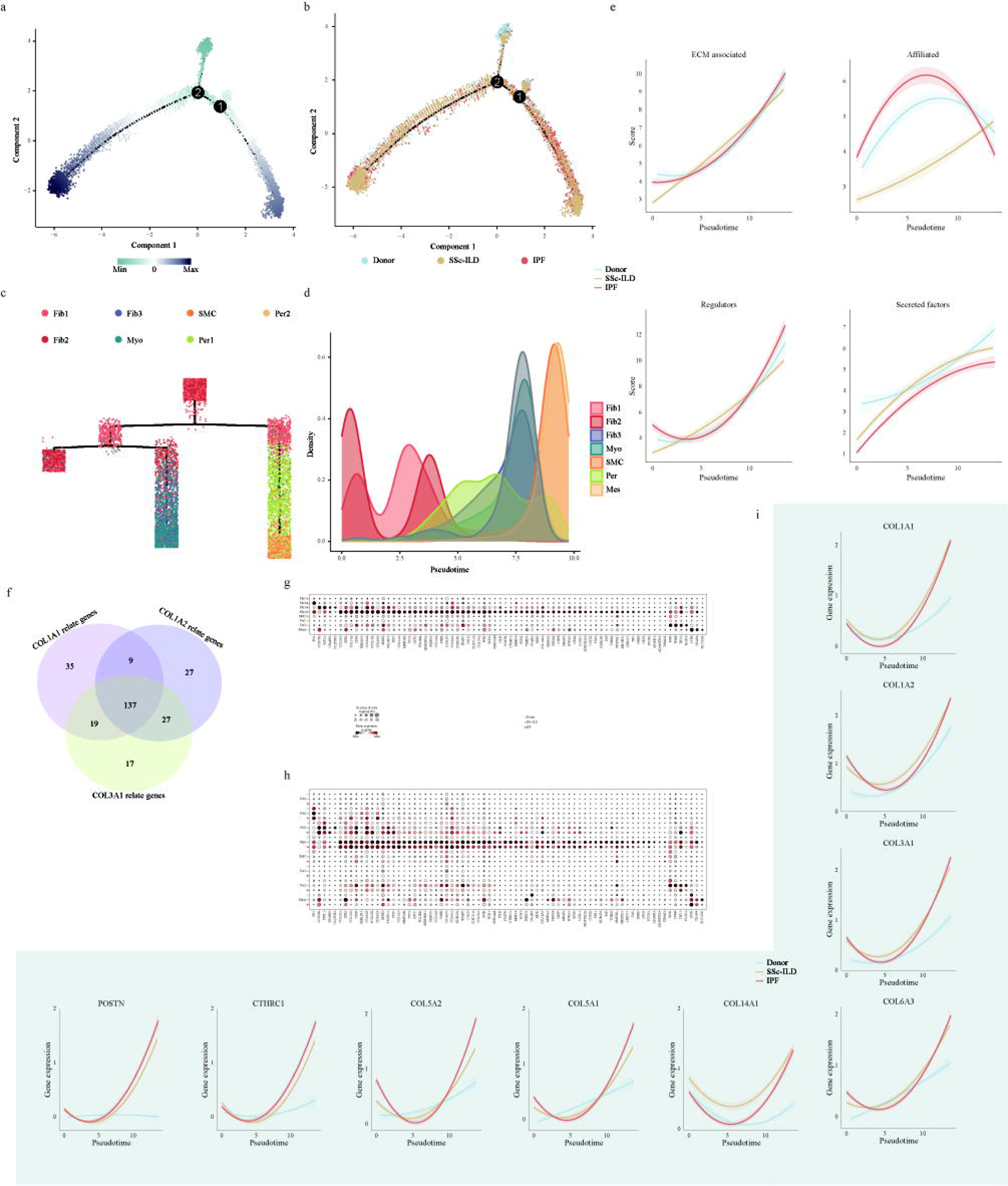
Trajectory analysis of mesenchymal cells and screening strategy for candidate genes. a-b, Trajectory analysis annotated by pseudotime and groups. c, Trajectory analysis annotated and overlaid with color by cell type class. d, Density plot of each mesenchymal subset along the pseudotime. e, Scores of ECM, collagens, proteoglycans, glycoproteins, ECM-associated genes, ECM-affiliated genes, regulators and secreted factors for each mesenchymal subset plotted along the pseudotime. f, Venn diagram showed the common genes of the top 200 genes, including positive and negative correlation of COL1A1, COL1A2, and COL3A1, respectively. g, Dot plot of relative expression of these common genes for each mesenchymal subset. h, Stratification of relative expression of these collagen genes from above correlate genes for mesenchymal subsets by groups. i, Candidate key genes expression in myofibroblasts stratified by groups plotted along the pseudotime.

**Fig. s6.**
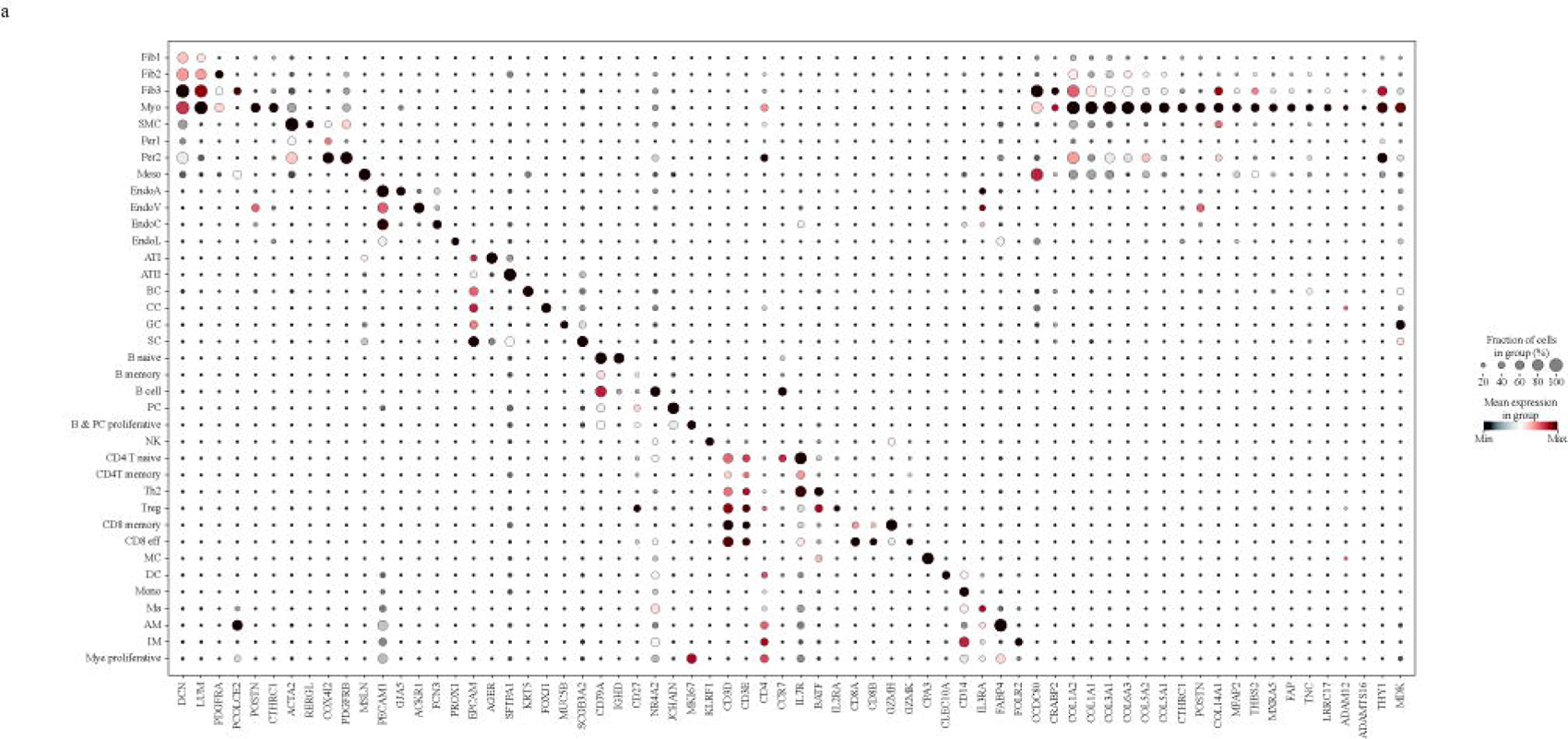
Relative expressions of 22 common genes in diversity cells of scRNA data. a, Dotplot showed relative expression of genes for all cells in scRNA data.

**Fig. s7.**
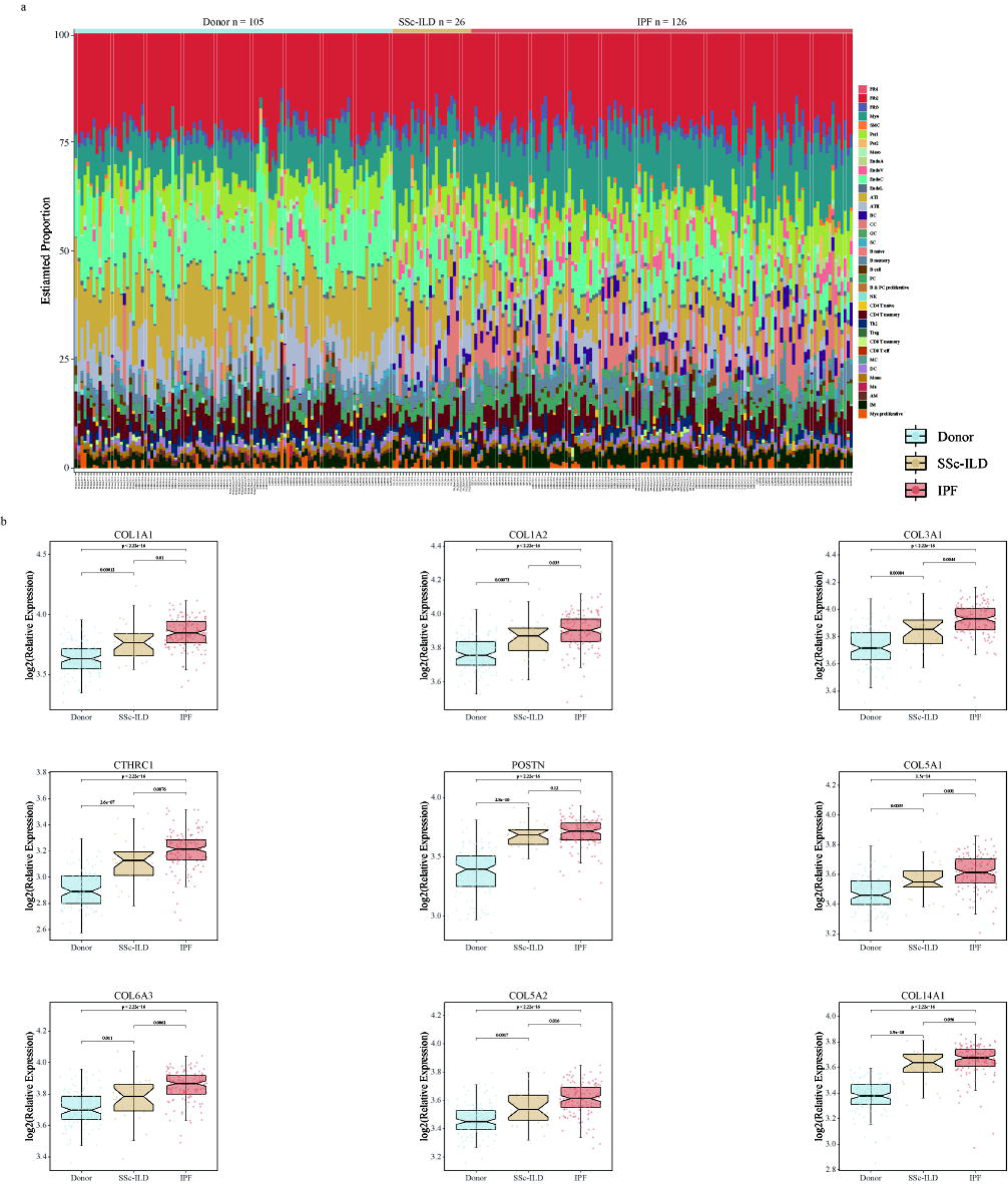
Ratio of myofibroblasts and Relative expressions of collagen genes in different bulk groups. a, Bar plot showed the estimated proportion of diverse cell types in each sample from bulk data. b, Box plot showed the relative expressions of nine collagens in the donor, SSc-ILD and IPF groups.

**Fig. s8.**
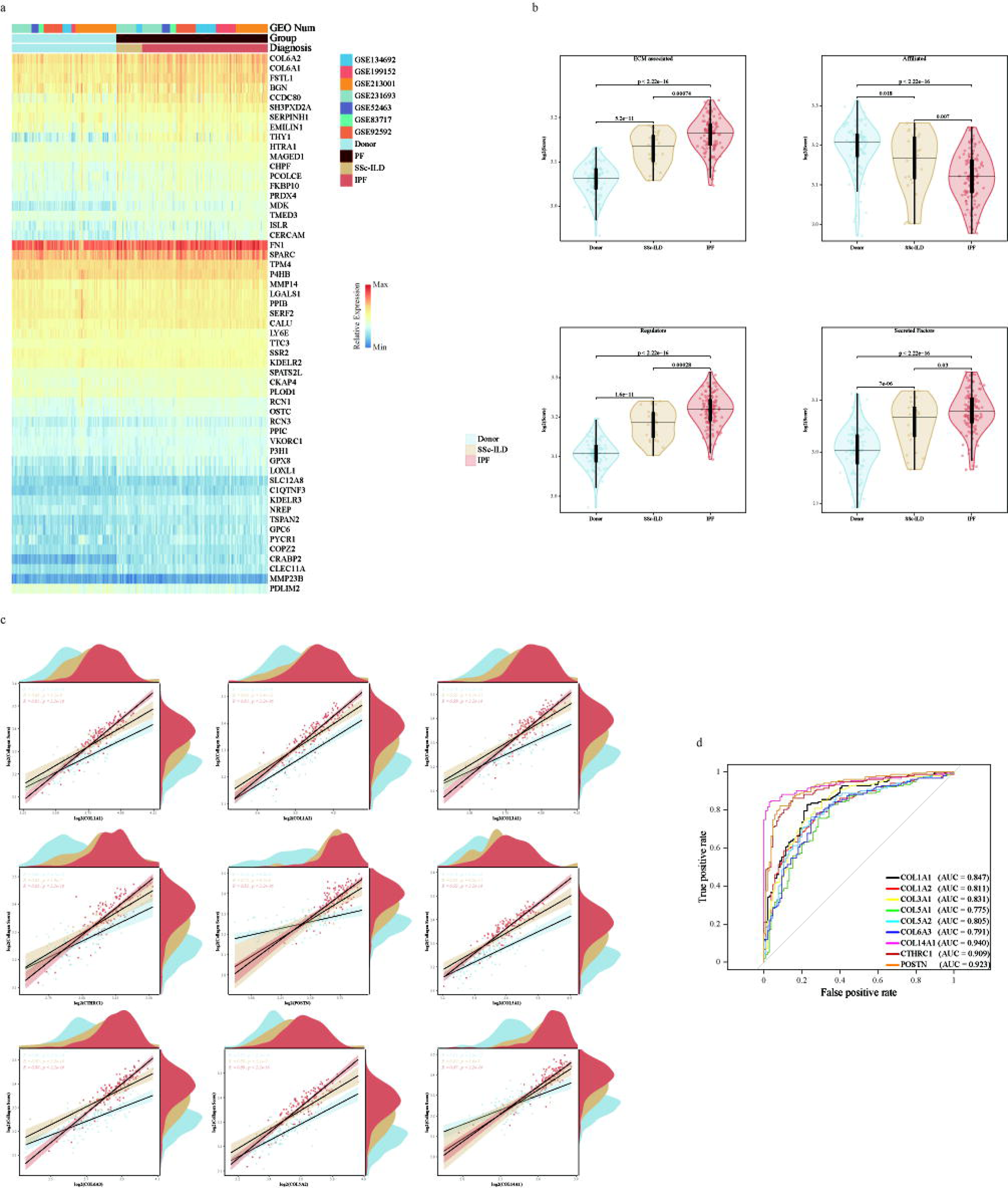
ECM associated scores and the correlation and ROC scores of nine collagens in different groups. a, Heatmap showed the expression of remaining genes in bulk RNA-seq results. b, Violin plot showed the scores of associated genes, ECM-affiliated genes, regulators and secreted factors for donor, SSc-ILD and IPF groups. c, The scatter plot showed a positive correlation between the nine collagen genes and the collagen score in the donor, SSc-ILC, and IPF groups. d, ROC plot showed favorable results for nine collagen genes in the PF and donor groups.

**Fig. s9.**
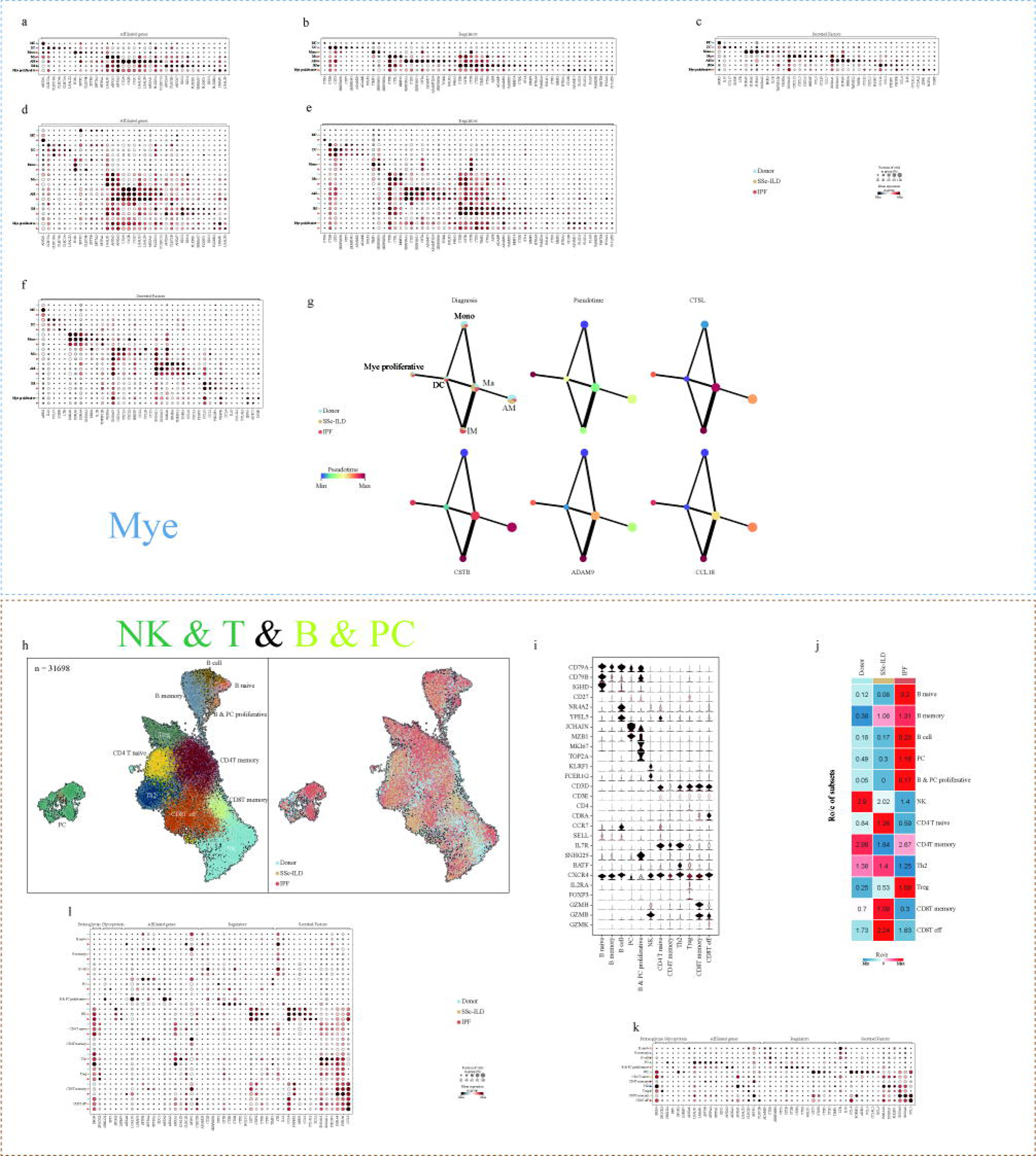
Immunological characteristics of myeloid subsets and lymphoid subsets in fibrosis microenvironment. a-c, Dot plot of relative expression of ECM affiliated genes, regulators and secreted factors for each myeloid subset. d-f, Stratification of relative expression of genes for myeloid subsets by groups. g, PAGA analysis of myeloid subsets stratified by groups, pseudotime and the hallmark genes of ECM regulators. Line thickness corresponds to the level of connectivity. h, UMAP embedding of 31,698 single cells in lymphoid subsets and stratification of cells by groups. Labels refer to the 12 subsets identified. B cell; B memory; B naïve; B &PC proliferative; CD4T memory; CD4 T naïve; Treg; Th2, T helper 2 cells; CD8T eff, CD8T effector cell; CD8T memory; CD8T effector cell; NK, natural killer cell; PC, plasma cell. i, Violin plot of expression of classical marker genes of each lymphoid subset. j, Group prevalence of each lymphoid subset estimated by Ro/e score. k, Dot plot of relative expression of proteoglycans, glycoproteins, ECM affiliated genes, regulators and secreted factors for each lymphoid subset. l, Dot plot of relative expression of genes for lymphoid subsets stratified by groups.

**Fig. s10.**
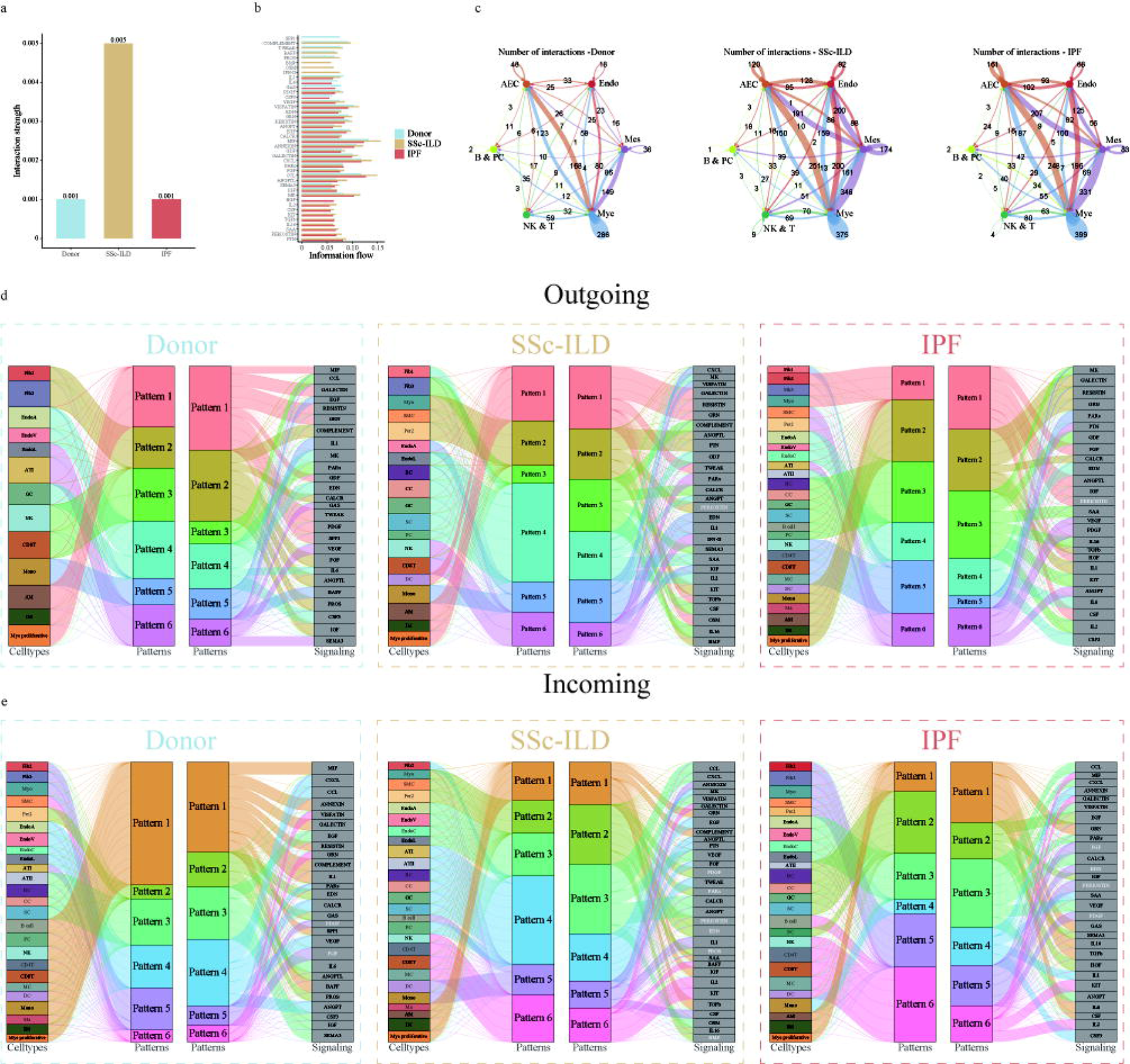
Identification of the ligand/receptor pair in PF. a, Interaction strength among Donor, SSc-ILD and IPF groups. b, The information flow of differentially expressed genes (DEG) for the PF stratified by groups. c, Number of interactions among six main cell types stratified by groups. d, The outgoing communication patterns of secreting cells stratified by groups. Key signaling was highlighted in white. e. The incoming communication patterns of target cells stratified by groups. Key signaling was highlighted in white.

